# Machine learning-inferred and energy landscape-guided analyses reveal kinetic determinants of CRISPR/Cas9 gene editing

**DOI:** 10.1101/2024.04.30.591525

**Authors:** Lei Jin, Risi Liyanage, Dongsheng Duan, Shi-Jie Chen

**Affiliations:** Department of Physics, Department of Biochemistry, and MU Institute for Data Science and Informatics, University of Missouri, Columbia MO 65211; Department of Molecular Microbiology and Immunology Department of Neurology, School of Medicine; Department of Biomedical Sciences, College of Veterinary Medicine; and Department of Bioengineering, University of Missouri, Columbia MO 65211, USA

**Keywords:** RNA folding kinetics, Folding energy landscape, CRISPR, spCas9, Cleavage efficiency, Off-target effects

## Abstract

The CRISPR/Cas nucleases system is widely considered the most important tool in genome engineering. However, current methods for predicting on/off-target effects and designing guide RNA (gRNA) rely on purely data-driven approaches or focus solely on the system’s thermal equilibrium properties. Nonetheless, experimental evidence suggests that the process is kinetically controlled rather than being in equilibrium. In this study, we utilized a vast amount of available data and combined random forest, a supervised ensemble learning algorithm, and free energy landscape analysis to investigate the kinetic pathways of R-loop formation in the CRISPR/Cas9 system and the intricate molecular interactions between DNA and the Cas9 RuvC and HNH domains. The study revealed (a) a novel three-state kinetic mechanism, (b) the unfolding of the activation state of the R-loop being the most crucial kinetic determinant and the key predictor for on- and off-target cleavage efficiencies, and (c) the nucleotides from positions +13 to +16 being the kinetically critical nucleotides. The results provide a biophysical rationale for the design of a kinetic strategy for enhancing CRISPR/Cas9 gene editing accuracy and efficiency.

## Introduction

CRISPR (clustered regularly interspaced short palindromic repeats)-Cas9 (CRISPR -associated protein 9) system, one of the most powerful tools in biological science [1], is widely employed in live-cell imaging [2, 3], gene engineering [4, 5], gene knockdown [6, 7], and gene therapy [8]. Associated with the (abut 100-nucleotide (nt)) single guide RNA (sgRNA) precisely targeting the DNA strand with 20-nt targeting recognition segment [1], the *Streptococcus pyogenes* (Sp) Cas9 binds to the double-stranded (ds) DNA target by recognizing the protospacer adjacent motif (PAM) and establishing base pairing between the guide RNA and the DNA target. Subsequently, SpCas9 endonuclease cleaves the dsDNA with its RuvC and HNH domains [9–11]. The complementary sequence match between the 20-nt target-recognition segment of the sgRNA and the target DNA strand constitutes an “on-target” cleavage site on the dsDNA [9–13]. Besides the on-target sites, partial complementary between sgRNA and target DNA sequences can also result in the binding of Cas9 and cleavage at such “off-target” sites [9–13]. Off-target cleavage can cause undesirable genetic alterations and should be avoided [14]. Maximizing the on-target efficiency and minimizing the off-target effect are significant challenges in developing the CRISPR-Cas9 system [15–23]. To optimize the CRISPR-Cas9 system, we need to identify the key determinants for the on/off-target cleavage efficiencies and, based on the physical mechanism, to develop computational tools to predict cleavage efficiencies and to design optimal sgRNA.

In recent years, bioinformatics or machine learning-based algorithms have been developed to evaluate on-target efficiency and identify potential off-target sites [24–45]. For instance, the in vivo library-on-library methodology sgRNA Scorer developed by Chari et al. [24] and sgRNA design rules Azimuth devised by Doench et al. [26] can be used to assess sgRNA on-target activity and help sgRNA selection; MIT (MIT Zhang’s model) [28], CCTop (CRISPR/Cas9 target online predictor) [29] and CFD (cutting frequency determination)[26] can be used to evaluate off-target cleavage potential. However, these data-driven approaches rely solely on the available experimental datasets and do not address the physical mechanisms underlying the cleavage. The free energy based physical approach uCRISPR [44] shows that on-target cleavage efficiency and off-target effects can be described in a unified framework by considering the folding stability of the R-loop complex. Although by accounting for the thermal equilibrium stability of the system, the physical model enables a better performance in evaluating the cleavage efficiency of both on- and off-target sites, the general Spearman rank correlation with the test set is below 0.50 and the predictions for off-target site are not always satisfactory.

Structural biologists have identified the essential structures in the cleavage reaction of the CRISPR-Cas9 system [46, 47] and a structure-based molecular mechanism of the cleavage reaction has been proposed [48], where the Cas9 protein recognizes sgRNA to form the Cas9-sgRNA complex, which then recognizes the PAM in the double-stranded DNA (dsDNA) target genome [49, 50]. The PAM for Cas9 is generally a 5’-NGG-3’ motif that can be recognized by the Cas9 through water-mediated hydrogen bonds in the minor groove of the dsDNA duplex [13, 47], and the binding of the Cas9-sgRNA complex to the dsDNA at the PAM site results in local destabilization of the adjacent DNA base pairs, initiating the unwinding of the target dsDNA from the PAM-proximal end [48]. If the downstream DNA sequence is complementary to the sgRNA spacer segment, the target strand of the unwound dsDNA hybridizes with the RNA strand, initiating a zipping-like reaction toward the PAM-distal end of the dsDNA. The process results in the formation of an R-loop with a 20-base pair (bp) RNA-DNA hybrid duplex and a single-stranded (ss) non-target DNA strand [51]; see Fig. 1A-C.

**Figure 1:**
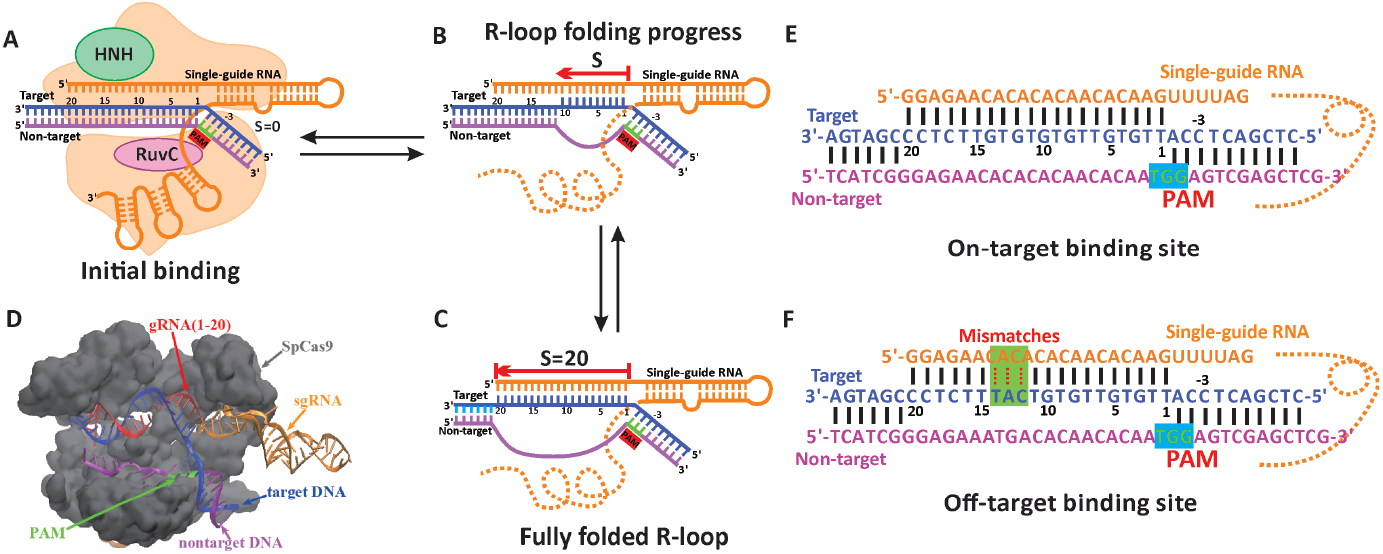
(A)-(C) The folding process of the R-loop. (D) The structural information of Cas9-sgRNA-dsDNA ternary complex. (E) On-target binding site: R-loop with complementary RNA-DNA hybrid. (F) Off-target binding site: R-loop with mismatched RNA-DNA hybrid.

In most previous theories, the system was assumed to be in thermal equilibrium, and cleavage occurred after the formation of the R-loop. Thus, the efficiency was tied to the equilibrium population of the R-loop [52]. However, the theories are not consistent with off-target cleavage data, where a less stable RNA-DNA duplex can lead to high cleavage efficiency. Recent kinetic experiments did not support a strong correlation between Cas9-sgRNA-dsDNA thermal equilibrium binding affinity and off-target cleavage efficiency, with distinct tolerance for mismatches in the PAM-proximal and -distal regions [48, 53, 54]. The results of these kinetic experiments suggest that cleavage efficiency is not necessarily correlated with binding equilibrium [53]. Furthermore, experimental evidence points to the kinetic control of the CRISPR cleavage activity, especially for off-target sites. However, current kinetic models [55, 56] are not accurate enough to quantitatively explain the kinetic mechanism.

In the present study, we leverage the large amount of cleavage data [7, 26, 28, 32, 40, 57], an ensemble machine learning algorithm [58, 59], kinetic simulation, and the rigorous free energy landscape analysis to explore the kinetic mechanism from which we develop an accurate predictive tool for CRISPR/Cas9 gene editing. Here are several essential features of the study:

1. Accounting for nucleotide-nucleotide and nucleotide-Cas9 interactions, using a random forest machine learning algorithm, we infer the kinetically critical determinants from which we construct zipping-like, step-by-step state transitions in the process of the formation and disruption of the R-loop. Kinetic Monte Carlo (KMC) simulation and free energy landscape analysis together predict the ensemble of trajectories and the transition rates between different states.
2. Large-scale simulations and kinetic profiling for the sequences in the different databases of on-/off-target cleavage reveal the active state (AS), the critical structures that initiate the cleavage induced by the RuvC and HNH domains of Cas9. We have identified the partially formed R-loop containing at least 17 consecutive base pairs (from the PAM site) as the cleavage-active state (AS). The result is different from the previously proposed fully formed R-loop (containing the full 20 base pairs) as the AS.
3. Moreover, the combined data and kinetic analysis uncovers a long-lived intermediate state: the R-loop with 10-12 consecutive base pairs (the seed region formed). The folding and unfolding kinetics of the R-loop (i.e. the targeting process) act as a two-step (three-state) reaction process.
4. To search for the kinetic determinants for the cleavage efficiency, we leverage the experimental dataset, the random forest ensemble learning algorithm, and various thermodynamic and kinetic features of the system, and have found the overall unfolding rate of the AS as the key determinant that shows the strongest correlation with the experimentally measured cleavage efficiency data. The unfolding rate of the AS, which captures the key kinetic feature of the cleavage process, accurately predicts the on-target activity and off-target effect of a CRISPR-Cas9 system with a higher accuracy than other models. Moreover, this kinetic model may serve as a new, advanced tool for rational design of optimal sgRNA for a given target.

## Results and Discussion

### A data-driven learning for the kinetic and thermodynamic features suggests that the R-loop unfolding rate is the most influential factor for cleavage efficiency

Using the uCRISPR energy parameters [44], we performed KMC simulations to obtain the kinetic information for the R-loop folding/unfolding for all the 14538 cases in the training dataset, see Table S1 for details. From the simulated trajectories and the Vfold model for nucleic acids folding [60], we considered a large set (*>* 10) of relevant kinetic and thermodynamic features as candidates of the determining features for CRISPR efficiency. These candidate features include the folding free energy Δ*G*_R*−*loop_, the equilibrium population 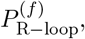, and the folding 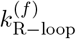 and unfolding 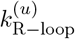 rates of the R-loop; see Fig. 2A for a set of 10 selected feature variables.

**Figure 2:**
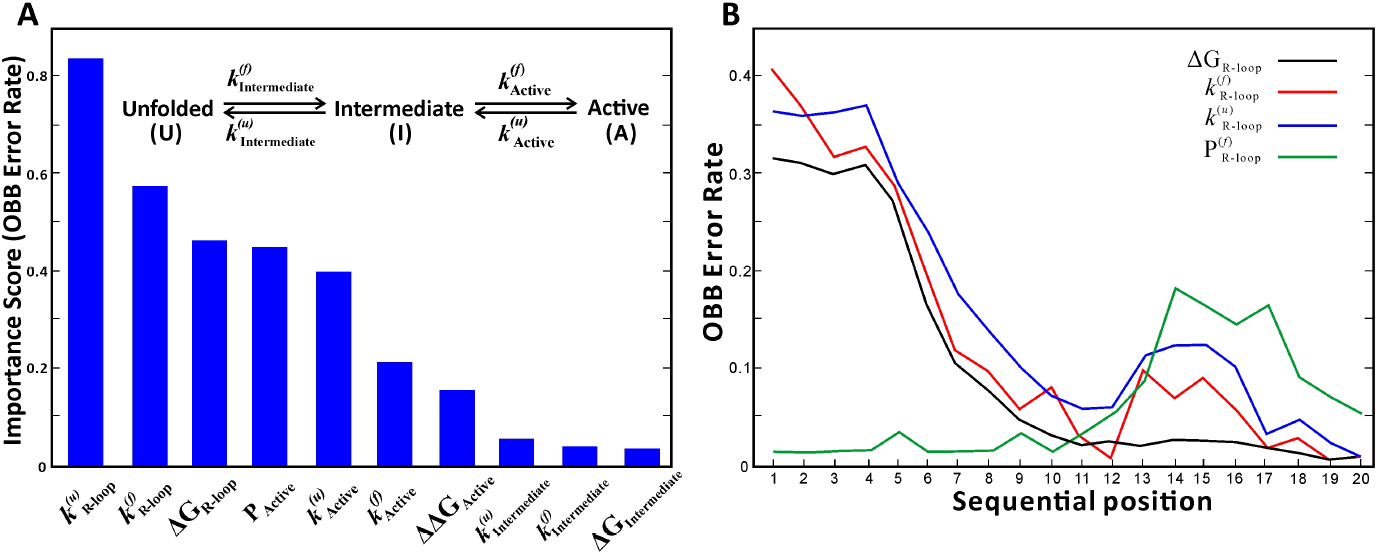
(A) Ranked importance score (OOB Error Rate (OER)) for 10 candidate feature variables in the kinetic model. Here 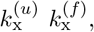, and Δ*G*_x_, and are the unfolding rate, the folding rate, and the folding free energy of state “x”. *P*_active_ is the equilibrium population of the AS. ΔΔ*G*_active_ is the free energy of the AS relative to the intermediate state. The intermediate state and the AS are defined based on the KMC simulation trajectories and folding free energy landscape. See details in the section “Two-steps reaction mechanism” (B) The calculated OOB Error Rate in the random forest classification algorithm for the different sequential positions for various feature variables.

Here, the thermal equilibrium properties are calculated from the Vfold model [60] and the kinetic-related properties are extracted from KMC simulations. Our purpose here is to identify the most important feature variables that determine the CRISPR-Cas9 cleavage efficiency.

The CRISPR cleavage efficiency data from experiments for a given target is in the form of *η*([*S*]) = {[*S*], *η*}, where [*S*] = {*S*_*target*_, *s*_1_, *s*_2_, …, *s*_20_} denotes the target DNA sequence *S*_*target*_ and the guide RNA sequence with *s*_*n*_ being the nucleotide at positions +*n* (*n* = 1, 2, …, 20). To evaluate the importance of each feature variable, we convert the original dataset into the form of *η*([*F*]) = {[*F*], *η*}, where [*F*] = {*f*_1_, *f*_2_, …, *f*_*m*_} denotes the *m* feature variables (see the x-axis labels of Fig. 2A for *m* = 10 feature variables) with *f*_*l*_ (*l* = 1, 2, …, *m*) being the *l*-th feature variable for the given sequence *S*. We use Random Forest (RF) to evaluate the importance of each feature variable. RF is a supervised machine-learning algorithm based on an ensemble of decision trees, where each tree is a configuration of different series of decisions each leading to an outcome of the cleavage efficiency. The RF trains the decision trees such that for a given input of feature variables, the RF gives the cleavage efficiency with the least ambiguity. Here RF is used for its ability to sample a wide range of data for training, overcome bias and overfitting in the learning process, and account for the different combinations of input features [58, 59] through the use of random subspace method (feature bootstrap aggregation).

The learning power of a decision tree can be measured by its test results on the out-of-bag (OOB) data set, i.e., the data not included in the training set for the learning of the decision tree. The OOB error for a decision tree is OOB-averaged difference between the decision tree-predicted and the experimentally determined cleavage efficiency. The average OOB error (OE) is the OOB error averaged over the ensemble of the decision trees. As shown below, the OE can provide measure for the importance of the feature variables. Specifically, to determine the importance of a feature variable *f*_*k*_, we randomly shuffle its values between the different data sets and use the changed data as the input for the RF to predict the cleavage efficiency and compute the OE. The feature variable *f*_*l*_ that shows the greatest change in OE is the most critical feature for CRISPR efficiency; see the section “Random Forest approach” in Supplementary Information (SI) for details. As shown in Fig. 2A, the importance scores (normalized OE, namely OOB error rate, OER) for the 10 candidate feature variables suggest that the R-loop unfolding rate 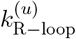 is the most important feature that determines the cleavage efficiency.

### Machine learning identifies the key nucleotides that impact CRISPR cleavage efficiency

#### The critical nucleotides

RF and the experimental data have identified the R-loop unfolding rate as the kinetic determinant for cleavage efficiency. To pinpoint the most crucial nucleotides for cleavage efficiency, we further employ RF to identify the positions (from +1 to +20) of the nucleotide sequence that most critically impact the R-loop unfolding rate. Our data set for the decision trees consists of data points in the form of 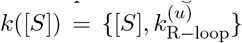 for each target sequence [*S*]. The nucleotides at the different positions impact the R-loop unfolding rate through their interaction energies. To examine the importance of the nucleotides at the different positions, we convert [*S*] into the energy parameters: 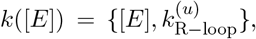, where [*E*] = {*E*_1_, *E*_2_, …, *E*_20_} denotes the (position-dependent) energy parameter with *E*_*n*_ being the energy parameter for the nucleotide at position +*n*. Physically, *E*_*n*_ reflects the free energy change for the disruption of the DNA base pair at position *n* followed by the formation of the base pair between the gRNA and the target DNA nucleotide (at position *n*) and the potential contact between the non-target DNA nucleotide (at position *n*) and the Cas9 protein. The 14538 different sequences from 11 individual experiments form the ensemble of data points for the learning of RF. See Table S1 for dataset details To identify the most important nucleotide positions, we compute the aforementioned OER for each position *n* = 1, 2, …, 20 by randomly replacing its energy parameter *E*_*n*_ with the ones of the same position in other data sets.

As shown in Fig. 2B, the OER for the equilibrium folding stability Δ*G*_*R−loop*_ of the R-loop shows three phases: a slow decrease from +1 to +5, a steep decrease from +6 to +10, and a plateau from +11 to +20. The result indicates a monotonically decreasing importance of nucleotide identity for the R-loop folding stability as the position moves away from the PAM site. In contrast, the non-equilibrium unfolding rate of the 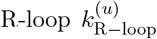 shows a drastically different behavior: the OER decreases continually to a low trough in the seed region [+1, +10] then rises to a relatively high plateau from +13 to +16 before decreasing from +17 to +20. The result suggests the critical importance of nucleotides from +13 to +16 for the R-loop unfolding kinetic rate and hence the CRISPR-Cas9 cleavage efficiency.

#### Two distinct regions

Fig. 2B for the position-dependent OER profile suggests two important regions for determining the cleavage efficiency: the seed region from +1 to +10 and the kinetically important distal region from +13 to +16. Previous studies have demonstrated a direct correlation between the cleavage efficiency and the thermal equilibrium stability of the seed region [53]. In contrast, mutational experiments have shown that the cleavage efficiency lacks a direct correlation with the equilibrium thermal stability of the distal region, especially for the kinetically important region (+13 to +16) [53]. The result suggests that the seed region can quickly reach thermal equilibrium while the [+13, +16] region may not reach equilibrium during the target recognition process, therefore, it is important to consider the non-equilibrium kinetics (for the R-loop unfolding).

### Energy landscape analysis and kinetic simulation suggest a three-state kinetics of the R-loop

#### On- and off-target energy landscapes

For an off-target binding, the non-Watson-Crick (mismatched) base pairs between the sgRNA and the DNA target are less stable than the corresponding Watson-Crick (canonical) base pairs. As a result, the energy landacpe of an off-target sequence is elevated by the mismatches and often shows bumps at the positions where mismatches occur (Figs. 3 and 4). The distorted energy landscapes for off-target cases may lead to distinct kinetics of R-loop folding unfolding and hence the kinetic progression of the CRISPR cleavage.

**Figure 3:**
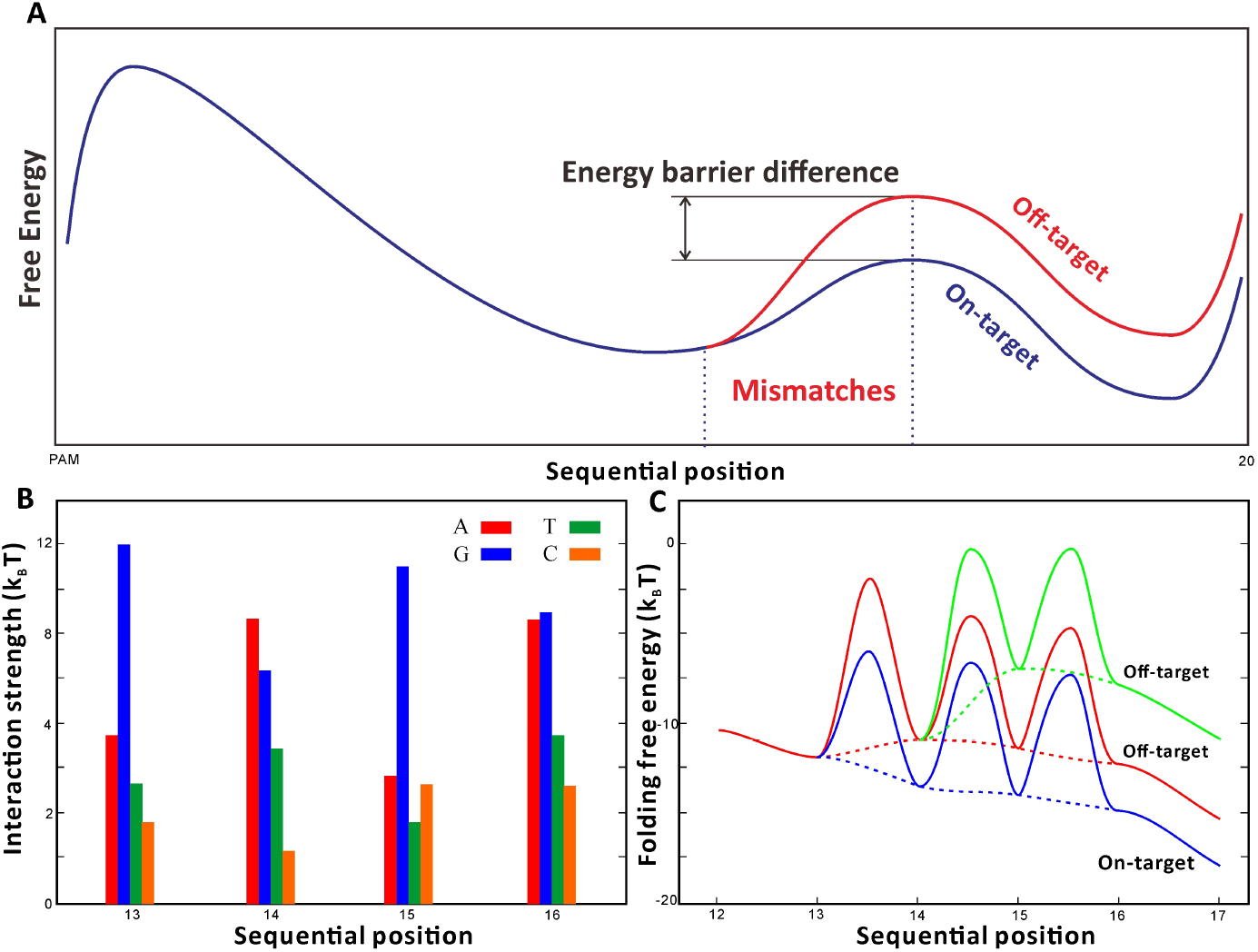
(A) The different free energy landscapes for on-target and off-target cleavages. The off-target sequence (red line) contains mismatches at positions 12-14, and compared to the corresponding on-target sequence (blue line), an off-target sequnce shows an increased energy barrier for the transition from the partially folded to the fully folded R-loop. (B) The interaction strength between the non-target DNA strand and the Cas9 RuvC domain extracted from molecule dynamics simulations. (C) The overall (dashed lines) and zoom-in (solid lines) folding free energy landscape for an on-target sequence (EXM1 target 1 [4], blue lines) and off-target sequences with a single mismatch at positions +14 (red lines) and +15 (green lines), respectively.

**Figure 4:**
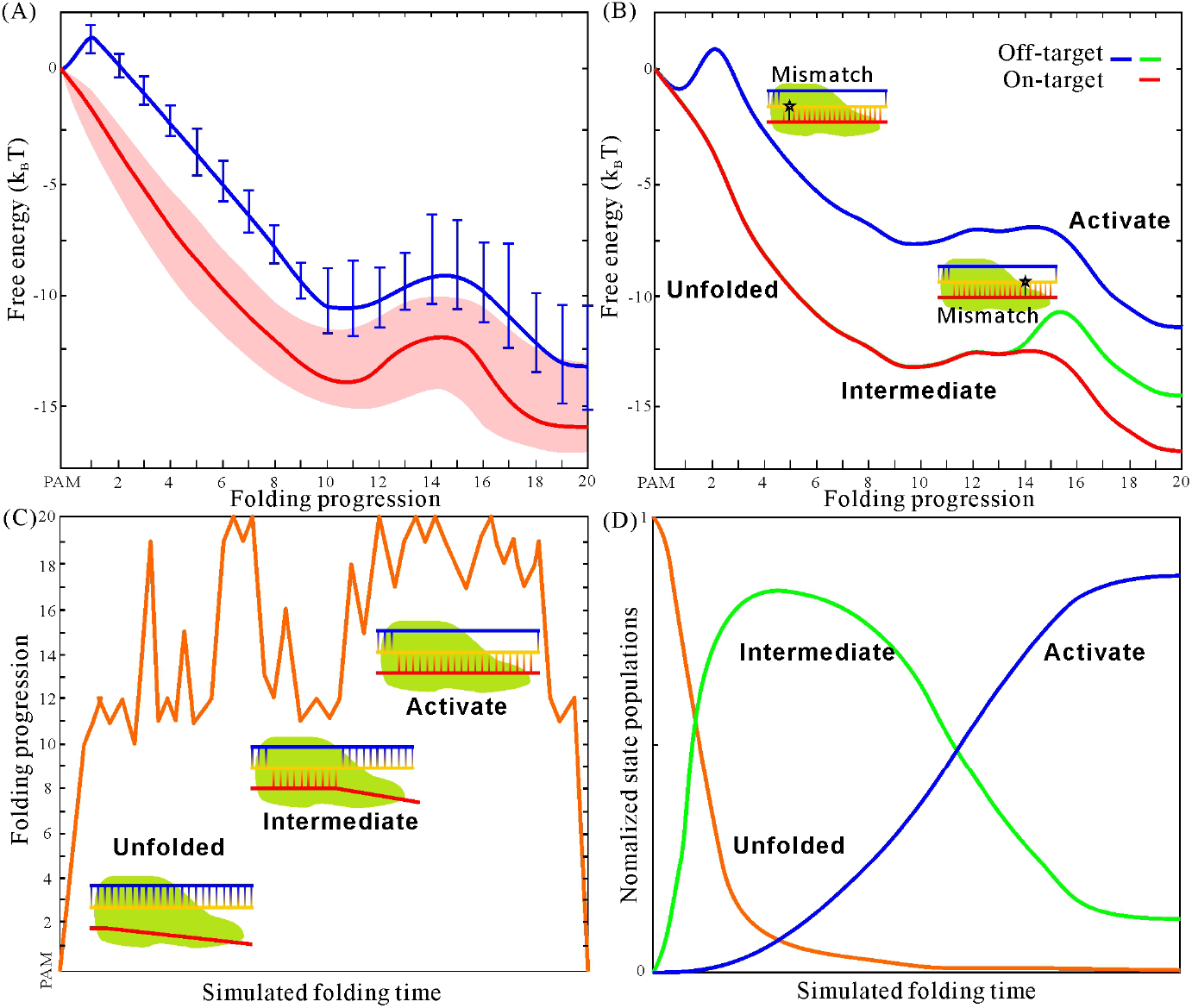
(A) The calculated free energy landscapes of the R-loop folding for the on-target EXM1 dataset [4]. The red line and the pink shadow represent the average free energy profile and the landscape fluctuations for the different sequences in the dataset [4], respectively. The blue line shows the average single mismatch free energy penalty (free energy shift of the R-loop caused by the single mismatch) at different positions and the error bars demonstrate the distribution of the free energy penalty of different mismatches at each position. (B) Predicted free energy landscape of the R-loop for the on-target sequence EXM1 target 1 [4] (red line) and the corresponding off-target caess with a single mismatch at PAM-proximal (blue line) and PAM-distal (green line) regions. The intermediate state and the AS correspond to the two long lived states shown in the folding trajectories. (C) R-loop folding trajectory of an on-target sequence (EXM1 target 1 [4]). Schematic diagrams illustrating the 2D structures along the folding trajectory are shown. (D) The simplified folding pathway with the unfolded state, the intermediate state, and the AS.

In general, off-target sequences have a lower folding stability and tend to have a faster unfolding rate of the R-loop, thus a lower cleavage efficiency. However, the high cleavage activity observed in certain off-target cases remains perplexing from a equilibrium thermodynamic point of view. To investigate the impact of the mismatches on CRISPR cleavage efficiency, we analyzed the overall folding free energy landscape of different on-target sequences and their corresponding off-target variants with single mismatches. Fig. 4A shows the averaged folding free energy landscape of the on-target EXM1 dataset [4] and the free energy penalties of the different mismatches at different positions. Clearly the folding free energy penalties (resulting in an upward shift of the off-target sequence R-loop folding free energy) for mismatches in the seed region can significantly increase the folding free energy landscape, placing the landscape far above the original on-target sequences. On the other hand, mismatches at the distal region may not elevate the folding free energy landscape high enough above the original on-target sequences. When considering the variation in the free energy landscape of on-target sequences (pink shadows in Fig. 4A) and the mismatch free energy penalties, We note that some off-target cases may exhibit a lower folding free energy than that of a on-target case (for a different site), Likewise, an on-target case can have a similar or even higher unfolding free energy barrier (i.e., a slower R-loop unfolding rate) than some off-target cases. These features can partially explain the mismatch tolerance at the distal region, making it understandable for some off-target sequences with mismatches at the distal region to exhibit relatively high cleavage efficiency. To fully comprehend the mechanism of on-target activity and off-target effects, the overall determinants of cleavage efficiency should be thoroughly investigated.

#### Three states

The KMC trajectories for the folding of a typical R-loop (e.g. EXM1 target 1 gene [4] in Fig. 4C) show three (long-lived) characteristic states: the unfolded state with 0-9 bp hybridization between the guide RNA and the target strand of the DNA; the partially folded structures with 10-16 bp hybridized base pairs, and the folded structures with 17-20 bp hybridized base pairs; see Fig. 4C.

The three states above are also shown as three distinct regions in the free energy landscape derived from the trajectory ensemble-averaged free energies (Fig. 4B). The landscape indicates an initial downhill folding for the formation of the first 10 base pairs in the seed region from positions 0 to 10, and after overcoming the kinetic barrier in the region from positions 13 to 16, a second downhill folding to complete the formation of the full 20 base pairs. The free energy landscape shows a local minimum (a metastable kinetic intermediate) corresponding to an ensemble of structures with 10-12 base pairs zipped up from the PAM end and a global minimum (fully folded R-loop) with around 20 base pairs zipped up from the PAM end. The result is consistent with the ensemble learning RF result about the critical role of the region from nucleotides +13 to +16: the macrostate of structures with 13-16 base pairs zipped up from the PAM end serves as the transition state (highest energy barrier) between the intermdiate state the folded R-loop state and hence determines the kinetic rate.

In most previously developed theoretical models, the fully folded R-loop structure is considered the functional structure for DNA cleavage. Through the analysis of folding free energy for different datasets, we have observed that the average free energy of the R-loop’s fully folded structure with all the 20 RNA-DNA hybrid base pairs formed is indistinguishable from the averaged folding free energy of partially folded R-loop structures with 17-19 RNA-DNA hybrid base pairs formed. Thus, the folded R-loop can be treated as a mixture of structures with 17-20 RNA-DNA hybrid base pairs formed. Experiments indicate that the sequence dependence of cleavage efficiency at distal positions 17-20 is weak for both on-target and off-target sequences. From this, we infer that structures with 17-19 RNA-DNA hybrid base pairs formed are equivalent in terms of cleavage activation. Under this assumption, an AS can be defined as a structure ensemble with 17-20 RNA-DNA hybrid base pairs formed.

In consistency with the long-lived structures observed in the kinetic folding trajectories, we define three macrostates: the unfolded state U, the intermediate state *F*_int_, and the AS *F*_act_, representing the structure ensemble with structures containing 0-9, 10-16, and 17-20 RNA-DNA hybrid base pairs, respectively. These three states are interconnected by the kinetic processes of R-loop formation and disruption. Consequently, the populational kinetics of the R-loop kinetic folding can be simplified from Fig. S1B to Fig. 4D.

### Two-step reaction mechanism predicts the cleavage efficiency kinetics

Based on the definition of the three states, the unfolded state “U”, the intermediate state “I”, and the AS “A”, we can established the overall folding kinetics as a two-step folding/unfolding reaction:

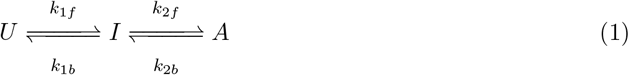

For most cases of the CRISPR-Cas9 systems, we found *k*_1*f*_ ≫ *k*_2*f*_ and *k*_2*b*_ ≫ *k*_1*b*_, suggesting a preequilibration between U and I for the folding and between I and A for the unfolding. Thus we have the following results for the overall folding *k*_*f*_ and unfolding *k*_*u*_ rates of the AS:

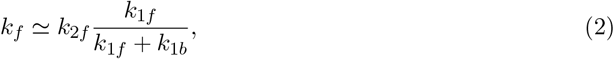

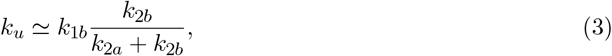

Where *k*_*f*_ and *k*_*u*_ are the overall folding and unfolding rates of the AS. *k*_1*f*_, *k*_2*f*_, *k*_1*b*_ and *k*_2*b*_ can be obtained from kinetic Monte Carlo simulations from Eq.6 and Eq.7 described in the Method section.

To validate the two-step folding/unfolding reactions model illustrated in Fig. 5A, we first examined the the AS by simulating the R-loop folding/unfolding kinetics for an on-target cleavage case and its corresponding off-target cases with various single mismatches. In these simulations, a cleavage state (where the dsDNA is cut by the Cas9) was added to the kinetics with a one-way transition rate *k*_cut_ from the AS to the final cleavage state. The dsDNA is cleaved upon entering the cleavage state, and the sgRNA-Cas9-DNA complex is completely deformed in the cleavage state. The total population of non-cleaved states, including the unfolded state, the intermediate state, and the AS, was calculated over time and shown in Fig. 5B and Fig. 5C. The consistency of the predicted population kinetics of the non-cleaved states with experimental measurements for both on-target and off-target sequences indicates that the definition of the AS effectively describes the cleavage-active reaction. Following this approach, facilitated by the knowledge of the cleavage reaction rate *k*_cu*t*_, we can simulate the full cleavage kinetics for both on-target and off-target sequences, and thereby evaluate on-target activity as well as off-target effects.

**Figure 5:**
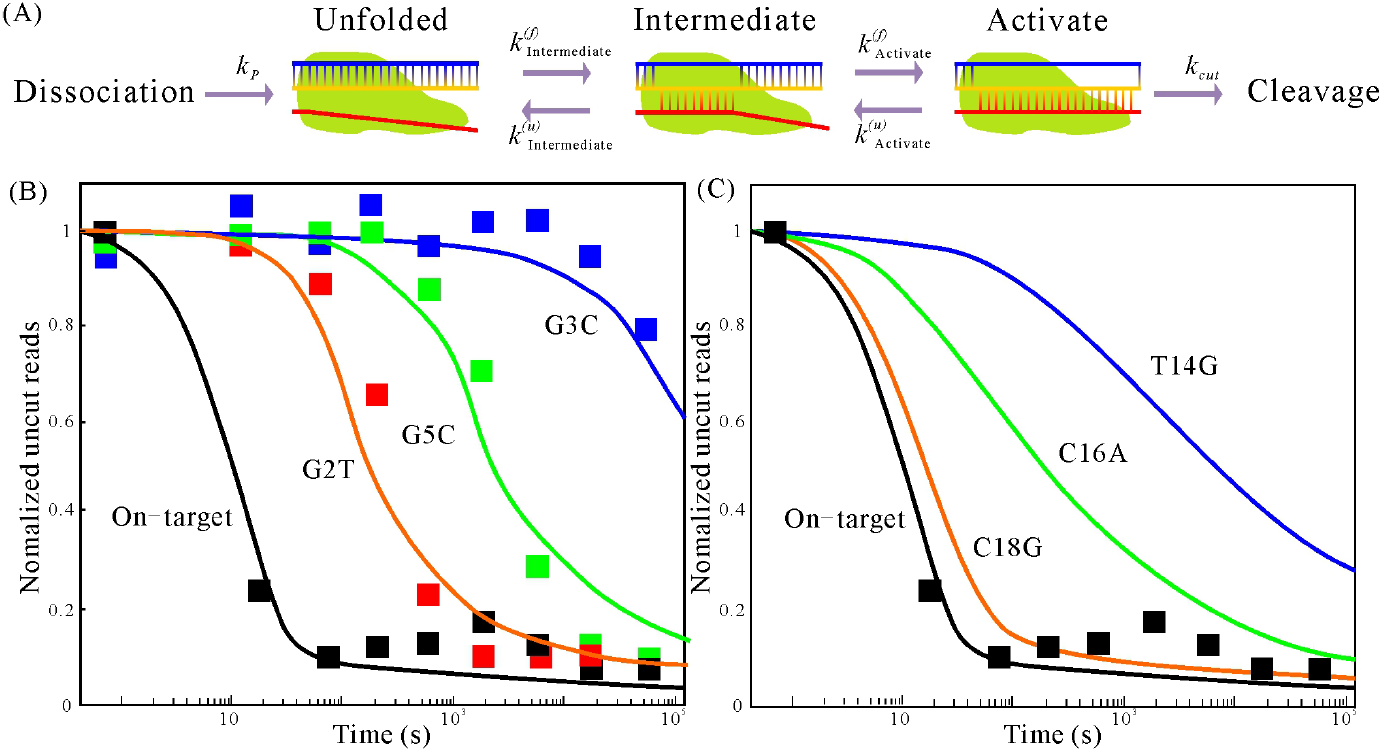
(A) A schematic illustrating the simplified kinetics of Cas9 cleavage efficacy, including Cas9-sgRNA-PAM binding, R-loop formation, and cleavage. (B) and (C) The simulated cleavage fraction of an on-target sequence (20-GACGCATAAAGATGAGACGC-1) and its corresponding off-target sites with single mismatchs in the PAM-proximal region (B) and PAM-distal region (C) as a function of reaction time. The symbols (squares) represent the experimentally measured data. [53].

### Kinetic model significantly improves off-target predictions

Through the RF supervised ensemble learning from the experimental cleavage data, we have identified the R-loop unfolding rate *k*_*u*_ (Eq.3) as the key determinant and hence an effective scoring function of the CRISPR cleavage efficiency. To further improve the accuracy of the *k*_*u*_-based cleavage prediction, we re-train (refine) the scaling factors of the energy parameters in 10 by repeating GA procedures. See details in “Method” under the supervision of the correlation between *ln*(*k*_*u*_) and experimental cleavage efficiency. After the parameter refinement, we re-examined the conclusion about the dominant role of the R-loop unfolding rate in determining the cleavage efficiency. We re-calculated the correlation between the experimental cleavage efficiency and the R-loop folding free energy Δ*G* and the folding *k*_*f*_ and the unfolding *k*_*u*_ rates of the AS for each data set. As shown in Fig. S3 for the Koike-Yusa/Xu Mouse ESC on-target dataset [33] as an example, *ln*(*k*_*u*_) shows a much stronger correlation with the experimental efficiency than Δ*G*. For 6 test data sets (3 on-target and 3 off-target), as shown in Fig. 6, *ln*(*k*_*u*_) shows the strongest correlation (by both the Spearman and the Pearson correlation coefficients) with the cleavage efficiency.

**Figure 6:**
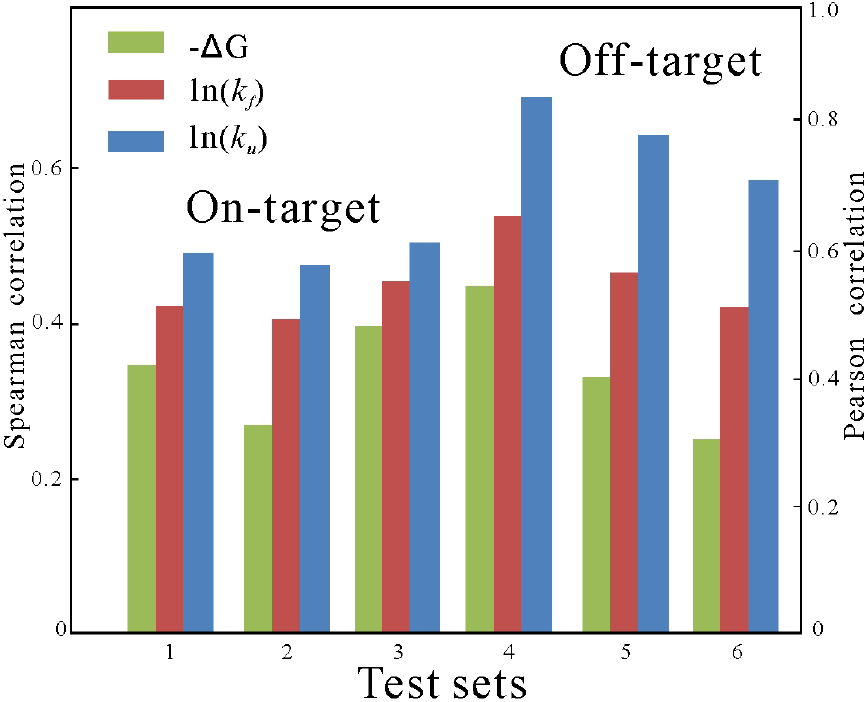
Overview of the correlation between the experimentally measured cleavage efficiencies and the calculated folding free energy, folding and unfolding rates for various types of datasets. 1-3 denote the on-target datasets (1, Koike-Yusa/Xu Mouse ESC [32]; 2, Hart Rpe [37]; 3, Hart Hct116-lib1 [37]) and 4-6 denote off-target datasets (4, Doench-CD33 part [7]; 5, Tsai-VEGFA site 2 [40]; 6, Kuscu-sgRNA 6 [33]).

The conclusion is consistent with that drawn from the random forest machine learning model. To further confirm the conclusion, we employed the re-trained (optimized) parameters to re-run the random forest feature importance analyses, (see the first subsection of “Result” and the section “Random Forest Approach” in SI)The results confirmed that the unfolding rates *k*_*u*_ obtained from Eq.3 are ranked as the highest important feature (Fig. S4 for details). Moreover, the kinetic properties as the top-ranked important features, as well as the higher correlations of the kinetic features (compared to the equilibrium thermodynamic features Δ*G* further support the importance of the kinetic mechanism to determine the CRISPR-Cas9 cleavage efficiency and the unfolding rate of the activating state *k*_*u*_ as the key determinant.

The overall average Spearman/Pearson correlations for 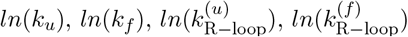 and ΔG for on-target/off-target training sets before and after the two rounds of parameter optimization are shown in Table 1. From Table 1, we can see that after parameter optimizations, the overall averaged Spearman/Pearson correlations for *ln*(*k*_*u*_) for both on-target and off-target sequences have improved. However, the improvements in correlation for on-target sequences and off-target sequences are different. Specifically, the correlation for on-target sequences has slightly increased by 0.04 (from 0.39 to 0.43), not as pronounced as that of the off-target sequences (increased by 0.08). We infer that the two-step reaction mechanism involving detailed structure transition in the distal region (+13 to +16) is more important in off-target cases, while on-target cases are essentially dominated by the seed region and are less affected by the distal region because the folding free energy landscape of on-target sequences is smoother compared to off-target sequences. The absence of a high energy barrier between the intermediate state and AS favors on-target sequences staying in the AS instead of the intermediate state. As a result, the R-loop unfolding rates are dominated by the seed region. Therefore, parameter optimization focused on the distal region (+13 to +16) for on-target sequences is not as important as for off-target sequences.

**Table 1:** correlation between feature variables and cleavage efficiencies. Spearman/Pearson correlations are calculated in the on-target/off-target training sets. The highest correlations for each parameter set are **bolted** and the overall highest correlations are **bolted**.

| OP | Initial<sup>a</sup> | Optimization 1<sup>b</sup> | Optimization 2<sup>c</sup> | Optimization 3<sup>d</sup> |
| --- | --- | --- | --- | --- |
| $\ln(k_u)$ | 0.36/0.74 | **0.39/0.77** | **0.43/0.85** | 0.41/**0.91** |
| $\ln(k_f)$ | **0.37/0.72** | 0.38/0.74 | 0.39/0.81 | 0.38/0.82 |
| $\ln(k_{R-loop}^{(u)})$ | **0.37/0.80** | **0.39/0.82** | 0.39/0.81 | 0.39/0.82 |
| $\ln(k_{R-loop}^{(f)})$ | **0.37/0.79** | 0.37/0.80 | 0.37/0.80 | 0.36/0.79 |
| $\Delta G$ | 0.33/0.75 | 0.33/0.72 | 0.33/0.73 | 0.33/0.76 |

<sup>a</sup> The initialized parameters directly obtained from singular value decomposition.
<sup>b</sup> Parameters trained with the correlation between $\ln(k_{R-loop}^{(u)})$ and experimental measurements.
<sup>c</sup> Parameters trained with the correlation between $\ln(k_u)$ and experimental measurements.
<sup>d</sup> Parameters trained with the correlation between $\ln(k_u)$ and experimental measurements only in the off-target training sets.

Considering the impact of the PAM sequence on the cleavage efficiency [26], we use the following scoring function for the cleavage efficiency:

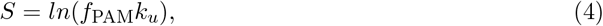

where the unfolding rate *k*_*u*_ is calculated by Eq.3 and *f*_PAM_ represents the effects of non-canonical PAM sequences (5’-NNN-3’) on the cleavage efficiency [26]. Since absolute cleavage efficiency is also significantly influenced by experimental conditions, and accounting for such effects is currently challenging in the scoring approach, as a result, the scoring function is more meaningful for the scoring of the different sequences/targets for a given system.

We first assessed the kinetic scoring approach on six on-target datasets, each containing more than 900 sgRNA sequences to maintain statistical significance. We evaluated the scoring results by calculating the Spearman correlation between the scores and the experimentally measured cleavage efficiency. Our results were compared to two other bioinformatics-based models, sgRNA Scorer [24] and Azimuth [26], as well as the thermodynamics-based model uCRISPR [44], as shown in Fig. 7A.

**Figure 7:**
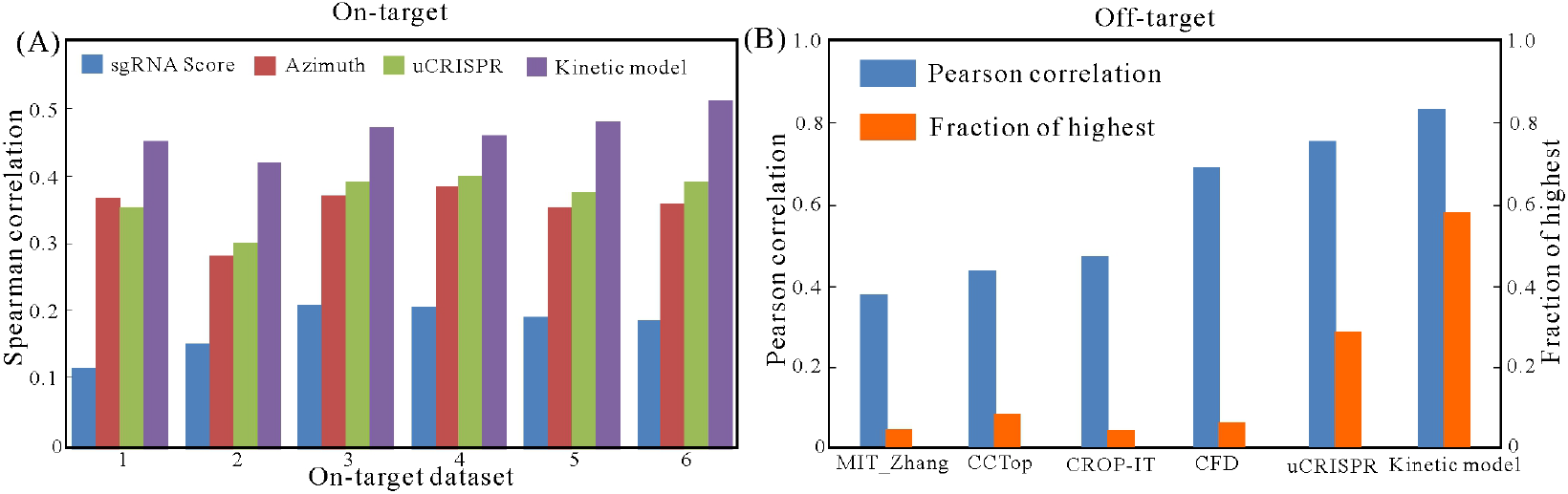
(A) Performance of on-target activity predictions compared with sgRNA Score [24], Azimuth [26] and uCRISPR [44]. 1-6 datasets are Koike-Yusa/Xu Mouse ESC [32], Hart Rpe [37], Hart Hct116-1 lib1 [37], Hart Hct116-2 lib1 [37], Hart HeLa lib 1 [37] and Hart HeLa lib 2 [37]. (B) Overall comparisons between different models with average correlation (blue bars) and fraction of best prediction with highest correlations (orange bars) for 51 off-target datasets.

For all six test datasets, the kinetic approach outperformed the other three models, achieving an average Spearman correlation of 0.47. In comparison, sgRNA Scorer, Azimuth, and uCRISPR yielded correlations of 0.18, 0.33, and 0.36, respectively. The introduction of kinetic mechanisms into the evaluation of on-target activity significantly improved the performance of model predictions, and in particular, the new kinetics-based predictions are more accurate than thermal stability-based predictions. To further validate the kinetic approach, additional datasets from different experiments were examined, and the results are shown in Supplementary Information Table S3.

We then test the model using off-target datasets. We collected and tested 51 independent experimental off-target datasets, which included both single and multiple mismatches across a wide range of genome-wide off-target sites. The Pearson correlation between the evaluated score and the experimentally measured cleavage efficiency was employed to assess the scoring results. Compared with four popular bioinformatics-based models and the uCRISPR model, the overall average Pearson correlation achieved by the kinetic model improved to 0.83±0.16, while the MIT Zhang [28], CCTop [29], CROP-IT [34], CFD [26] and uCRISPR [44] yield 0.38 ± 0.28, 0.43 ± 0.28, 0.47 ± 0.23, 0.68 ± 0.24 and 0.75 ±0.24, respectively (see Fig. 7B and the detailed results in Supplementary Information Table S4). Compared to uCRISPR [44], the number of failed cases in the current model (Pearson correlation *<* 0.4) was significantly reduced. This substantial improvement for the large number of sequences in datasets containing multiple mismatches indicates that the kinetic mechanism may indeed be a dominant factor that influences off-target cleavage efficiency.

## Conclusions

Since on- and off-target cleavage efficiencies vary across different sgRNAs, experimental conditions, and cellular environments, it is critically important to establish a physical mechanism-based fraemwork for optimizing sgRNA design and minimizing the off-target effects in the CRISPR-Cas9 cleavage system. Laveraging the large data for the cleavage efficiencies, by combinging supervised learning and physics-based kinetic modeling, we dentified the critical feature that determines the CRISPR/Cas9 efficiency for on- and off-target cleavages. Furthermore, we have developed a kinetic model that can demonstrate the step-by-step transitions of the R-loop folding/unfolding progression, allowing us to trace the kinetic trajectories of the system. By considering the non-target DNA strand interacting with the Cas9 RuvC/HNH domains at essential positions, the kinetic model, compared with other approaches, has acquired a significantly improved accuracy for the prediction of CRISPR/Cas9 cleavage efficiencies. Specifically, our comprehensive analyses on the large variety of sequences in the data bases of on- /off-target cleavage sites revealed the following detailed kinetic mechanisms in Cas9-sgRNA targeting and cleaving:

1. The folding and unfolding of the R-loop act as a two-step reaction process with a hybridized seed region formed as an intermediate.
2. Cleaving can be initiated after the AS with 17 continuous base pairs formed from the PAM end is formed.
3. The overall unfolding rate of the AS shows the most significant correlation with the experimentally measured cleavage efficiencies.

In contrast to the traditional thermodynamic analysis of the CRISPR/Cas9 system which assumes a two-state transition without considering the formation of the possible intermediates, the overall kinetics follows a two-step (three-state) reaction process. The first step in R-loop folding (seed region folding) essentially determines the binding affinity of Cas9-sgRNA to dsDNA, while the second step initiates cleavage as induced by RuvC and HNH domains. This mechanism may explain why binding affinity and cleavage efficiency weakly correlate in vitro and in vivo, especially for off-target sequences. Moreover, the strong tolerance of mismatches in the distal region suggests that the AS may contain multiple hybridized structures. The strongest correlation between states with at least 17 continuous base pairs formed in the R-loop and experimentally measured cleavage efficiencies verifies this mechanism.

## Methods

### R-loop folding kinetics

A functional CRISPR-Cas9 nuclease complex before the target DNA is cleavage contains the Cas9 bonded R-loop on the reaction channel between Cas9 RuvC and HNH domains (see Fig. 1D and Fig. S2 A). The single-stranded guide RNA’ hybridization with the 5’-*>*3’ target single-stranded (TS) DNA chain along the DNA unwinding enhance the affinity of the sgRNA-Cas9 binding to the target DNA duplex (see Figs. 1A-C). Since experiments suggest that the kinetic folding of R-loop have strong relation with the cleavage efficiency [48, 53, 54], we introduced the R-loop zipping-like kinetic folding/unfolding model following the nature of RNA/DNA double helical winding/unwinding (see Figs. 1A-C and Fig. S1 A).

### Folding states and kinetic moves

The formation of the R-loop is described as a zipping/unzipping-like base pair exchange between dsDNA and sgRNA-DNA hybrid from PAM-proximal to PAM-distal region (see Fig. 1A). The zipping pathway can be described with 20 states defined by the number of formed RNA-DNA hybride base pairs. Consistent with previous studies, the 10 bps of the R-loop adjacent to PAM form the seed region and the rest 10 bps belong to the PAM-distal region. One base pair switching from (to) dsDNA to (from) the sgRNA-DNA hybrid is defined as a basic kinetic move of the R-loop folding (unfolding). Specifically, a state of R-loop with N bps (namely state N) in hybrid has two kinetically adjacent R-loop states of N+1 bps and N-1 bps, respectively.

### Rate constants

We use the conventional Metropolis rule to calculate the rate constants of a kinetic move:

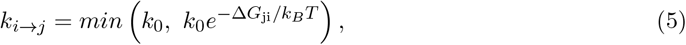

Where Δ*G*_ji_= *G*_*j*_ − *G*_*i*_ is the free energy different between the two states, *k*_0_ is the attempt frequency, *k*_*B*_ is the Boltzmann constant and T is the temperature. Therefore, the transition rates *k*_*f,n*_ from states N to N+1 for a folding step and *k*_*u,n*._ from states N+1 to N for an unfolding step are determined by the free energy landscape of the system.

### Kinetic Monte Carlo (KMC) simulation

The time evolution of the population for each state is described by the master equation 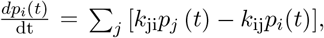, where *p*_*i*_ (*t*) and *p*_*j*_ (*t*) are the fractional population of state *i* and *j* at time *t*, respectively. The population kinetics, however, cannot provide information about the kinetic pathways. To reveal the microscopic trajectories of the R-loop folding, we employ kinetic Monte Carlo method [61] to simulate R-loop folding and unfolding pathways. Averaging over the ensemble of the simulated trajectories would yield the R-loop folding/unfolding pathway and the free energy landscape along the pathway.

In detail, a kinetic move in KMC is simulated in 3 steps:

1. Construct the total out-flow transition rate as *k*_total_ = ∑*k*_*p*_, where *k*_*p*_ is the rate constant for the transition from the current state *p* to one other adjacent state.
2. Select the next state *q* such that 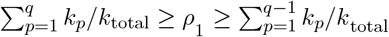 for a randomly generated number *ρ*_1_ between 0 and 1.
3. Update system time from t to *t* − *ln* (*ρ*_2_) */k*_total_, where *ρ*_2_ is another random number between 0 and 1.

Repeating the above kinetic moves starting from an initial state of the R-loop will generate a folding trajectory.

### Populational kinetics and overall transition rate

We perform multiple (e.g. 5000) KMC simulations to generate an ensemble of folding trajectories. Statistically, the average over a sufficiently large number of trajectories from the same initial state can give us the populational kinetics as well as the folding pathway; see Fig. S1B. More importantly, the overall transition rate from state A to state B can be calculated as

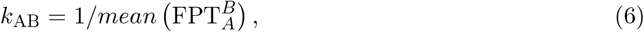

Where 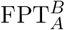 is the first passage time from A to B in a KMC trajectory. The corresponding reverse transition from B to A has the transition rate 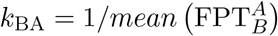 or

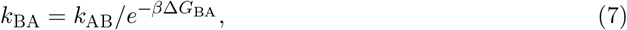

where Δ*G*_BA_ = *G*_*B*_ − *G*_*A*_ is the free energy difference between the two states. The free energy for a state such A and/or B that consists of an ensemble of microstates (structures) s is given by 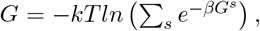, where *G*^*s*^ is the free energy of each microstate *s*.

### Energy parameterization

The free energy of state N with n base pairs in the RNA-DNA hybrid duplex from the PAM site is calculated as the sum of the base stacking free energies of the RNA-DNA hybrid:

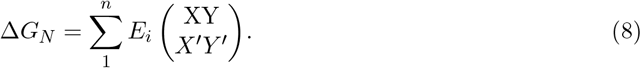

Where 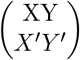 represents the base stack formed by the base pair 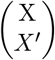 at position *i* and the base pair 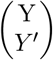 at position *i-*1. 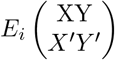 is the stacking energy, which is the free energy change in Eq.5 for the formation/disruption of a bae pair. There are 4^4^ = 256 energy parameters for all possible 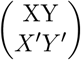 at each position of the sequence (1-20) including 4^2^ = 16 energy parameters for on-target sequences (canonical base pair stacking) and 240 energy parameters for off-target sequences (non-canonical base pair stacking). Considering the 20 positions of the R-loop, we have 20 x 16 = 320 and 20 x 240 = 4800 position-dependent energy parameters for the canonical and the non-canonical (mismatched) base stacks, respectively.

### Parameter optimization

We use the uCRISPR approach, which is based on the equilibrium thermal stability of the folded CRISPR/Cas9 system, to assign the initial values of the energy parameters in Eq.8. [44]. For th 4800 non-canonical parameters, to reduce the number of parameters, we decompose 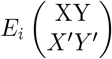 at position i:

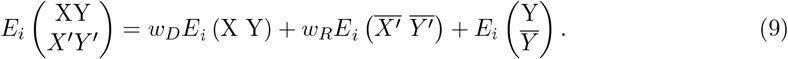

Where *E*_*i*_ (X Y) and 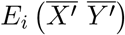 are the energy parameters for dinucleotides 5’XY3’ and 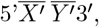 are *w*_*R*_ are the position independent weight coefficients, and 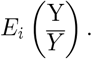. is the position-dependent non-canonical base pairing energy. 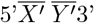 and 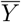 denote the dinucleotide complementary to 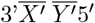 and the nucleotide complementary to Y, respectively. Accounting for all the non-canonical base pairing types, the energy term 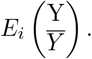. has 4 x 3 = 12 cases at the position of i. Therefore, I total, the above decomposition results in a reduction from 4800 to 242 parameters including the two position-independent weight coefficients.

The 320 sequence dependent energy parameters for the on-target sequences involved in this kinetic based model were obtained through singular value decomposition based on 5 training datasets: Wang/Xu HL60/KBM7 [32], Doench MOLM13/NB4/TF1 [7], Doench A375/AZD [26], Doench Mouse EL4 [7] and Koike-Yusa/Xu Mouse ESC [32], see Table S1 for details.

Similar to the on-target parameters the 242 off-target parameters can be determined using singular value decomposition with 18 off-target training sets, see Table S1 for details.

Before performing parameter optimization, we first use the Random Forest classification analyses [58, 59] to identify the most relevant kinetic features that determine the cleavage efficiency and the critical positions for the position-dependent energy parameters that need to be optimized. See SI for the details of the the Random Forest analyses.

As the last step of parameter optimization, the position-dependent energy parameters were sampled and optimized using the Genetic Algorithm between the non-target DNA strand and the Cas9 protein. See SI for the detailed optimizing procedure.

To obtain more detailed kinetic information in the R-loop folding/unfolding, we zoom into the detailed kinetic move the R-loop formation and deformation of the distal region. Different from the one step treatment of the RNA-DNA hybrid base pair formation for the seed region, we divided the RNA-DNA hybridization of the distal region into two parts: DNA base pair unwinding which involves the non-target DNA base binding to Cas9 protein, and target DNA base pairing with sgRNA’s corresponding base. Consequently, the free energy change of a single DNA-RNA hybrid base pair formation can be expanded as

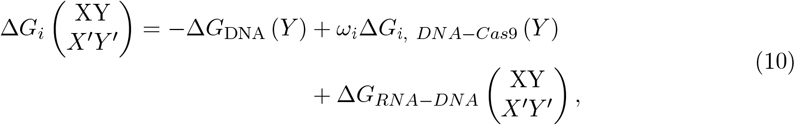

Where Δ*G*_*i, DNA*_ (*Y*) is the free energy of DNA base stacking at position i, 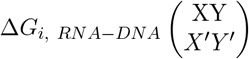 is the free energy of RNA-DNA hybrid base stacking at position i. Specifically, Δ*G*_*i, DNA−Cas*9_ (*Y*) explicitly describe the interactions between the non-target DNA base with the Cas9 protein. The free energy for all base stackings can be obtained through the nearest-neighbor model and related thermodynamic experiments [62]. The Δ*G*_*i, DNA−Cas*9_ (*Y*) should be physically determined without experimental information. And the position dependent scaling factors *ω*_*i*_ need to be trained to balance the composites.

### Molecular dynamic simulations

To determine the free energy of the interaction between the non-target DNA base and the Cas9 protein, we conducted molecular dynamic (MD) simulations to explore the detailed interactions between the DNA bases and the protein’s residues (refer to the supporting information for details). From these MD simulations, we extracted the interaction strength between the DNA bases and Cas9 HNH or RuvC domains at various positions in the sensitive distal region with different types of nucleotides, which were then used for calculating the free energy in Eq.10.

By conducting MD simulations, we observed a sequence-dependent, relatively strong interaction between non-target DNA and the Cas9 RuvC domain, as illustrated in Fig. 3B. We divided the kinetic movements into two parts and re-composed the free energy change (Eq.10) to calculate the state transition rate constants, as described previously. In Fig. 2C three examples are provided to depict the detailed folding free energy landscapes of an on-target sequence and two corresponding off-target sequences with single mismatches at +14 and +15 positions.

The interactions between non-target DNA and the Cas9 RuvC domain can either enhance or reduce the folding/unfolding rate of an off-target sequence with mismatches in the region of +13 to +16, depending on the sequences. Incorporating this detailed mechanism into the kinetic model should address the inconsistency observed in experiments regarding the correlations between mismatches and cleavage efficiency of off-target sequences.

### Genetic algorithm

According to Eq.10, there are four position-dependent scaling factors *ω*_*i*_ for the four essential positions distinguished by the Random Forest approach that need to be trained. We utilized a Genetic Algorithm (GA), a highly efficient simplex-like self-guided search approach, to train these parameters. The goal was to maximize the correlations between *k*_*u,R−loop*_ and experimentally measured cleavage efficiency in the training datasets, which were initially used for the initialization of energy

## Data availability statement

The data are available in the article and the online supplementary material.

