## Supplemental Information for "Machine learning-inferred and energy landscape-guided analyses reveal kinetic determinants of CRISPR/Cas9 gene editing"

### Energy and parameter frameworks

#### R-loop free energy calculation

Free energy of the state N with n base pairs in RNA-DNA hybrid along R-loop folding is calculated as the summation of the base stacking formation free energies change of the RNA-DNA hybrid:

$$\Delta G_N = \sum_1^n E_i \binom{XY}{X'Y'}, \tag{S1}$$

Where $\binom{XY}{X'Y'}$ represent the base pair $\binom{X}{X'}$at position *i* stacking with base pair $\binom{Y}{Y'}$ at position *i*-1. The free energy change between two kinetic connected states i and i-1 involved in Eq. 1 can be written as $\Delta G_{i(i-1)} = E_i \binom{XY}{X'Y'}$. For instance, $E_3 \binom{AG}{UC}$ represents the free energy of RNA-DNA hybrid base pair dG-rC at the 3rd position adding to the RNA-DNA hybrid helix with the dA-rU base pair at the 2nd position. The $E_i \binom{XY}{X'Y'}$ accounts for all the interactions in the CRISPR-Cas9 system including the free energy change of unwinding the DNA base pair stacking at position i and i+1, free energy change of RNA-DNA base pair stacking formation at position i-1 and i and free energy change of the RNA/DNA-Cas9 interaction rearrangements:

$$E_i \binom{XY}{X'Y'} = -\Delta G_{i,DNA} \binom{Y}{\bar{Y}} + \Delta G_{i,RNA-DNA} \binom{X}{X'} + \Delta G_{i,RNA/DNA-Cas9} \binom{XY}{X'Y'}. \tag{S2}$$

In Eq. S2, the $\Delta G_{i,DNA} \binom{Y}{\bar{Y}}$ and $\Delta G_{i,RNA-DNA} \binom{X}{X'}$ can be directly obtained through the Nearest-Neighbor model [2]. However, $\Delta G_{i,RNA/DNA-Cas9} \binom{XY}{X'Y'}$ is impossible to be exactly obtained for all the sequence cases. So, it is reasonable to composite the 3 energy terms in to an effective free energy change $E_i \binom{XY}{X'Y'}$ and fit for the effective value of $E_i \binom{XY}{X'Y'}$ among training data sets.

#### On-target energy parameters

For the on-target sequences, the nucleotide on the sgRNA at a sequential position is complimentary to the nucleotide on the target DNA at the same sequential position. $E_i \binom{X\,Y}{X'Y'}$ can be simplified as $E_i \binom{X\,Y}{\bar{X}\,\bar{Y}}$ or $E_i(X\,Y)$ and the total number of base stacking type at position i is 16. For the entire 20 positions, there are 320 parameters in total.

#### Off-target energy parameters

For the off-target sequence with single or multiple mismatches at position i, the free energy change of adding a new base pair stacking to the R-loop RNA-DNA hybrid helix $E_i \binom{XY}{X'Y'}$ can have 240 cases (without on-target cases). The total number of the parameters for the 20 positions is 4800 in total. To reduce the total number of the parameters, we calculate $E_i \binom{XY}{X'Y'}$ as

$$E_i \binom{XY}{X'Y'} = w_D E_i(X\,Y) + w_R E_i(\overline{X'}\,\overline{Y'}) + E_i \binom{Y}{\bar{Y}}, \tag{S3}$$

Where $E_i(X\,Y)$ and $E_i(\overline{X'}\,\overline{Y'})$ are the on-target energy parameters at position i, $w_D$ and $w_R$ are position independent weight constants, $E_i\binom{Y}{\bar{Y}}$ are position dependent non-canonical base pairing energy. Accounting for all the non-canonical base pairing types, the $E_i\binom{Y}{\bar{Y}}$ has 12 cases at the position of i. Therefore, there are 242 parameters in total for the entire 20 bps R-loop including 2 position independent weight constants.

**Parameter initialization**

Based on the previous study for the physical correlation between the R-loop folding free energy and the cleavage efficiency, the initial parameters were trained following the free energy orientated scoring method used in the uCRISPR model [1]. The 320 sequence dependent energy parameters for the on-target sequences involved in this kinetic based model were obtained through singular value decomposition with the involvement of 5 training datasets: Wang/Xu HL60/KBM7 [3], Doench MOLM13/NB4/TF1 [4], Doench A375/AZD [5], Doench Mouse EL4 [4] and Koike-Yusa/Xu Mouse ESC [3], see Table S1 for details.

Similar to the on-target parameters the 242 off-target parameters can be obtained through singular value decomposition with 18 off-target training sets, see Table S1 for details.

**Parameter optimization**

With the selection of the essential positions +13~+16 (see the following section ***Random Forest approach***), we decomposed the free energy change of a single DNA-RNA hybrid base pair formation as

$$\Delta G_i\binom{XY}{X'Y'} = -\Delta G_{DNA}(Y) + \omega_i \Delta G_{i,DNA-Cas9}(Y) + \Delta G_{RNA-DNA}\binom{XY}{X'Y'} \tag{S4}$$

Where $\Delta G_{i,DNA}(Y)$ is the free energy of DNA base stacking at position i , $\Delta G_{i,RNA-DNA}\binom{XY}{X'Y'}$ is the free energy of RNA-DNA hybrid base stacking at position i. Specifically, $\Delta G_{i,DNA-Cas9}(Y)$ explicitly describe the interactions between the non-target DNA base with the Cas9 protein. The free energy for all base stackings can be obtained through the nearest-neighbor model and related thermodynamic experiments [2]. The $\Delta G_{i,DNA-Cas9}(Y)$ should be physically determined without experimental information. And the position dependent scaling factors $\omega_i$ need to be trained to balance the composites.

To determine the interactions $\Delta G_{i,DNA-Cas9}(Y)$ between the non-target DNA bases with the Cas9 protein, we performed Molecular Dynamics (MD) simulations to estimate the interaction free energy, see the following section ***Molecule Dynamic simulations*** for details.

According to the Eq. S4, there are 4 scaling factors $\omega_i$ for the 4 essential positions distinguished by random forest approach need to be trained. We employed *Genetic algorithm* supervised by maximizing the correlation between the kinetic scoring (R-loop unfolding rate $K_{u,R-loop}$, see the following section ***Random Forest approach* for details**) and the experimental measured cleavage efficiency to train these parameters in the training sets which were used for the initialization of the energy parameters.

**Random Forest approach**

Random Forest (RF) classification approach was applied for ranking parameter sensitiveness for scoring order parameter selection and essential sequential positions screening for parameter optimization [6,7]. RF programming via ALGLIB 3.16.0 (https://www.alglib.net/) and Breiman and Cutler's Random Forests for Classification and Regression v 4.6-14 (https://www.stat.berkeley.edu/~breiman/RandomForests).

*Order parameter selection*

Calculate OOB Error Rate for order parameters in R-loop folding/unfolding (including but not limited in free energy change $\Delta G_{R-loop}$, R-loop folding rate $K_{f,R-loop}$, R-loop unfolding rate $K_{u,R-loop}$ and folded state's equilibrium population $P_{f,R-loop}$).

a) Random Forest training for data set $\mathcal{D}_n = \{OP_k(i), Efficiency_{Expt}(i)\}_{i=1}^{n}$

   (n is the total number of samples, $OP_k$ is the $k^{th}$ order parameters considered in, $Efficiency_{Expt}(i)$ is the experimental measured cleavage efficiency for $i^{th}$ sample)

   i. Bootstrap aggregation [7]
      1. Construct subsets through random sampling with replacement
      2. Regression trees training in subsets
      3. Average trained regression trees
   ii. Calculate averaged Out-of-bag error over the forest for each data point during training

b) Variable importance measurement for feature k
   i. Permutated values of feature k among the data points
   ii. Calculate out-of-bag error for each data point after permutated
   iii. Calculate importance score by averaging difference in out-of-bag error before and after permutated (OOB error Rate).

c) Calculate importance score for all features and rank the variable by importance score

This procedure is to figure out the correlation strength between the kinetic/thermodynamic order parameters and the overall cleavage efficiency of the sequences. The order parameters with higher ranking can be used as the scoring function of the cleavage efficiency.

*Essential sequential positions screening*

Distinguish essential sequential positions for energy parameters

a) Random Forest training for data set $\mathcal{D}_n = \{E(i,j)_{j=1}^{20}, OP(i)\}_{i=1}^{n}$

   (n is the total number of samples, $E(i,j)$ is the applied energy parameters of $i^{th}$ data point at position j, $OP(i)$ is the value of the selected order parameters for $i^{th}$ data point)

   i. Bootstrap aggregation
      1. Construct subsets through random sampling with replacement
      2. Regression tree training in subsets
      3. Average trained regression trees
   ii. Calculate averaged Out-of-bag error over the forest for each data point during training

b) Energy parameter importance measurement for position j

i. Permutated values of energy parameters at position j among the data points
ii. Calculate out-of-bag error for each data point after permutated
iii. Calculate importance score by averaging difference in out-of-bag error before and after permutated (OOB error Rate).

c) Calculate importance score for all positions and rank the positions by importance score.

This procedure is to evaluate the sensitiveness of the order parameters to position-dependent energy parameters. Because the position-dependent energy parameters are the basic trainable parameters in this model, the order parameter with higher sensitiveness to the position-dependent energy parameters can better account for the sequence specificity.

**Molecule Dynamic simulations**

We choose the Cryo-EM structure of SpCas9-sgRNA-DNA ternary complex (PBD: 5y36) including non-target DNA strand as the initial pre-cut 3D structure of the system [1]. The MD simulations are designed to estimate the strength of the interaction between non-target DNA strand and Cas9 RuvC domain. The crystal structures without the full length non-target DNA strand are not considered.

To reduce to simulation complexity, the simulated structure contains R-loop hybrid, dsDNA with PAM, non-target DNA strand, Cas9 HNH domain and RuvC domain, see Fig S2B

*MD simulations*

Periodic simulation cells are constructed with the water shell of 12 Å. Hydrogen atoms were added assuming standard bond lengths and were constrained to their equilibrium position with the SHAKE algorithm. Salt concentration was considered as 0.08 mM of NaCl, in agreement with related experimental conditions.

Using the Amber ff12SB force field with ff99bsc0 correction [2] for DNA and ff99bsc0+χOL3 correction [3,4] for RNA and Åqvist force field [5] for Mg2+ together with TIP3P model for water [6], all the simulations were performed with GPU version of Amber 16. Long-range electrostatic interactions are evaluated using the smooth particle mesh Ewald (PME) method [7]. Temperature of the system was controlled by Langevin Dynamics, Pressure was controlled by coupling the system to Berendsen barostat. All simulations were performed as following protocol:

i. Energy minimization to relax the water and counter ions, RNA, DNA and proteins were fixed with harmonic position restraints of $300\ \mathrm{kcal/mol \cdot Å^2}$ .
ii. Heat up the system from 0 to 310K by 3 steps with in total 500 ps canonical ensemble (NVT) simulations with the RNA, DNA and proteins fixed.
iii. Perform 500 ps isothermal-isobaric ensemble (NPT) simulation of equilibration with the R-loop hybrid, dsDNA and proteins fixed while non-target DNA unfixed.
iv. Perform 50 ns NVT simulation for energy calculation with the R-loop hybrid, dsDNA and proteins fixed while non-target DNA unfixed.

To account for the different type of nucleotide at different sequential position, the nucleotides were manually mutated to cover all possible cases.

*Calculate free energy*

To calculate free energy of the interaction between non-target DNA nucleotide and RuvC $\Delta G_{i,DNA-RuvC}(Y)$, we performed Molecular Mechanics / Poisson Boltzmann Surface Area (MMPBSA) calculations [8] to estimate the binding free energy of non-target DNA strand based on the 50 ns MD simulations. The binding free energy of non-target DNA nucleotide was calculated as the binding free energy difference between the binding free energy of non-target DNA strand with and without typical nucleotide

**Prediction evaluations**

*Measurement metrics*

The Spearman rank correlations (equals to the Pearson correlation coefficient) between predicted scores and the experimental measured cleavage efficiency are calculated as

$$R_s = 1 - \frac{6\sum d_i^2}{n(n^2-1)}. \qquad S5$$

Where $d_i$ is the difference between the ranks of the predicted scores and experimental measured cleavage efficiency and n is the number of the sequence data cases. Predictions for on-target sequences are evaluated by the Spearman rank correlations for the presented kinetic model and other models used for comparison.

The Pearson correlations between are predicted scores and the experimental measured cleavage efficiency are calculated as

$$R_p = \frac{\sum(S_i-\bar{S})(Eff_i-\overline{Eff})}{\sqrt{\sum(S_i-\bar{S})^2}\sqrt{\sum(Eff_i-\overline{Eff})^2}} \qquad S6$$

Where $S_i$ and $Eff_i$ are the predicted score and experimental measured cleavage efficiency for the $i^{th}$ sequence in the dataset, respectively. $\bar{S}$ and $\overline{Eff}$ are the averaged predicted score and experimental measured cleavage efficiency for the datasets, respectively.

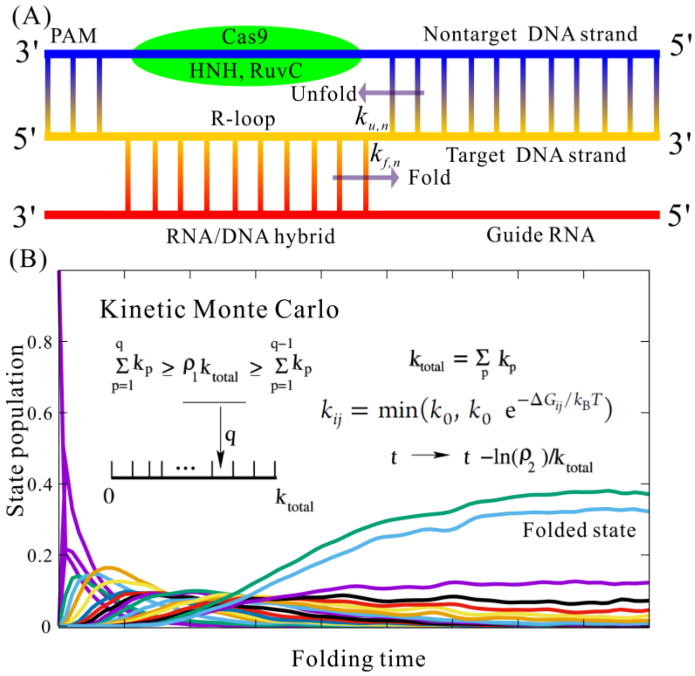


Figure S1. (A) The kinetic scheme of the kinetic model demonstrates the Cas9 cleavage efficacy. (B) Basic Kinetic Monte Carlo algorithm involved in the kinetic model and the folding pathway calculated through the ensemble of simulated folding trajectories.

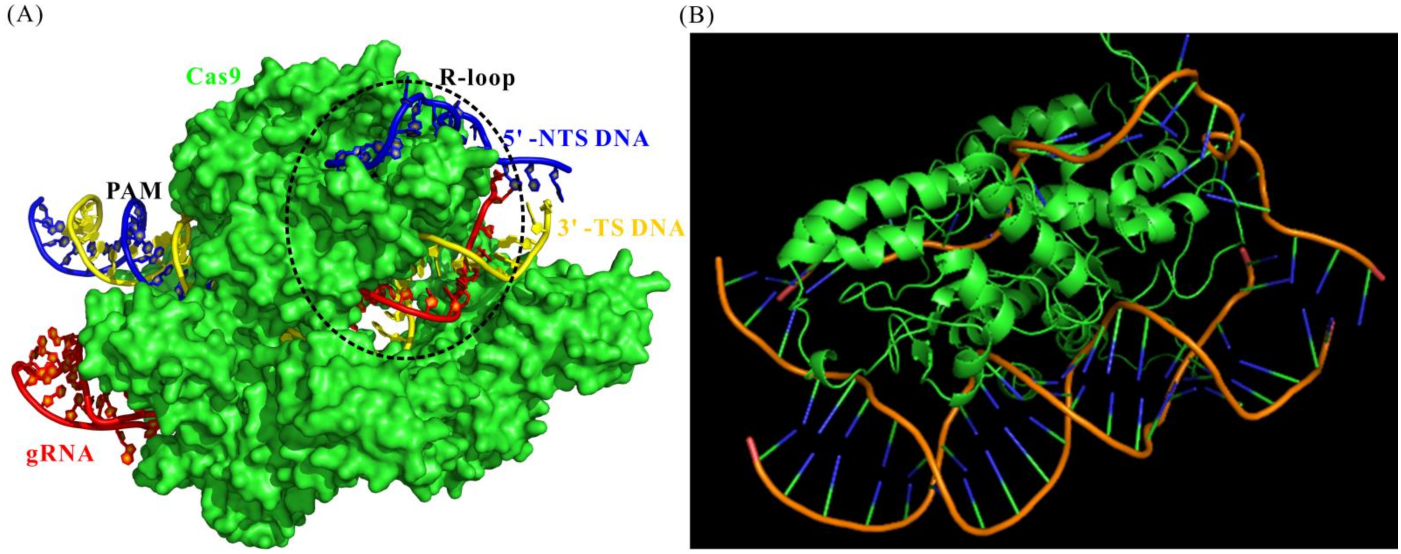


Figure S2. (A) The structural information of Cas9-sgRNA-dsDNA ternary complex with important factors considered in the kinetic model labeled on. The cycled region shows the essential interactions between the non-target single-strand (NTS) DNA and RuvC/HNH domains of Cas9. (B) Reduced MD simulation region focused on interactions between the non-target single-strand (NTS) DNA and RuvC/HNH domains of Cas9.

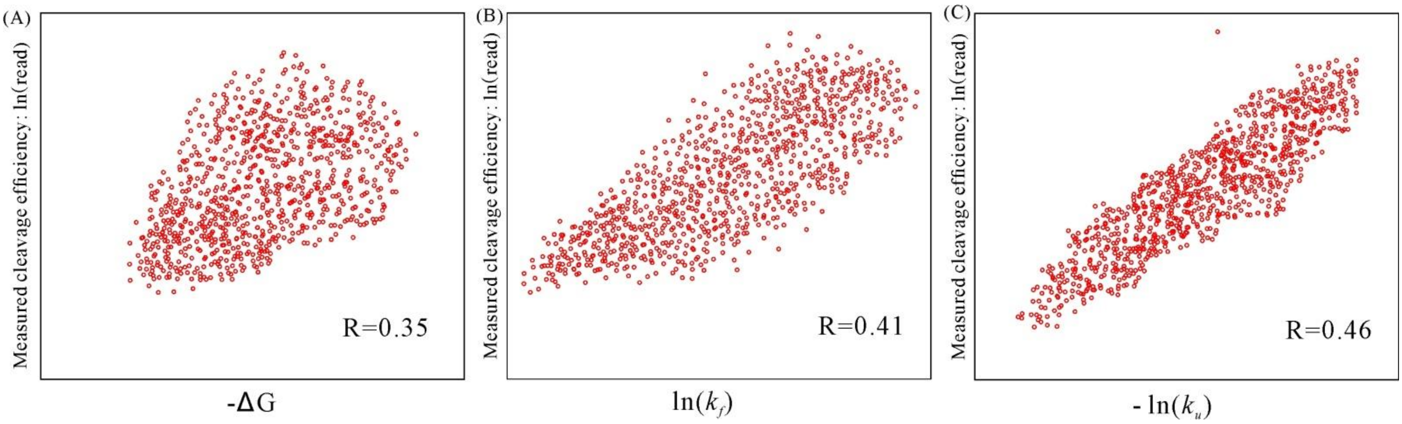


Figure S3. The correlation between the experimentally measured cleavage efficiencies and the calculated folding free energy (A), the folding rate (B) and the unfolding rate (C) for the 907 cases in the Koike-Yusa/Xu Mouse ESC on-target dataset [3], respectively. The calculated spearman correlations are labeled in each panel.

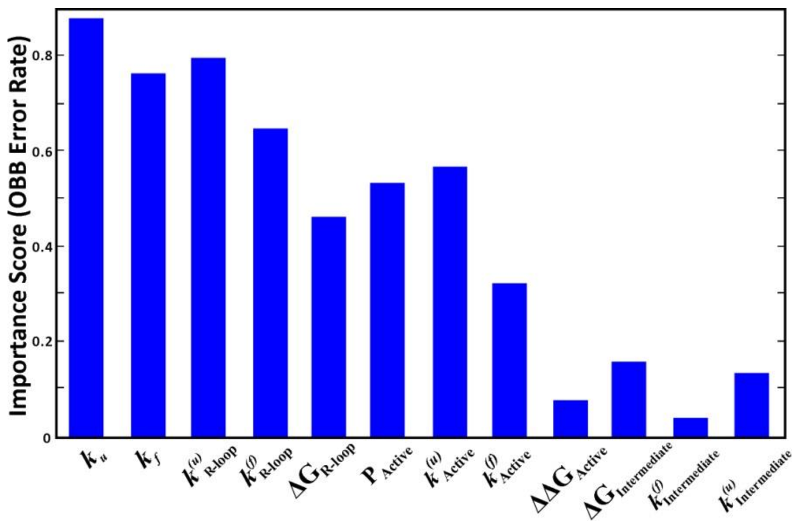


Figure S4. Ranked importance score (OOB Error Rate (OER)) for 12 candidate feature variables in the kinetic model after parameter optimization. Here $k_x^{(u)}$ , $k_x^{(f)}$ and $\Delta G_x$are the unfolding rate, the folding rate, and the folding free energy of state ``x''. $P_{\text{Active}}$ is the equilibrium population of active state. $\Delta\Delta G_{Active}$ is the free energy of the active state relative to the intermediate state.

**Table S1, Summary of datasets used in the kinetic model for parameter training.** On-target datasets are highlighted in light blue and off-target datasets are highlighted in light green. In total, 14538 data points were involved in the training data sets, including 7148 data points (sgRNAs) for on-target training and 7390 data points (off-target sites) for off-target training. Datasets Kuscu-sgRNA 1-12 are only used in parameter optimization while other datasets are used both in parameter initiation and optimization. Usage notes generally describe the characteristics of the data sets that support the usage in model developing.

| Dataset | Ref. | Number of gRNAs/sites | Species & Biology | Efficiency Quantification | Usage notes |
|---|---|---|---|---|---|
| **Wang/Xu HL60/KBM7** | 3 | 2076 | Human HL60/KBM7 cells | log2 fold-change | On-target data sets with various gRNAs for human. Consistent quantification method. Frequently-used for model developing. |
| **Doench MOLM13/NB4/TF1** | 4 | 881 | Human MOLM13/NB4/TF1 cells | log2 fold-change | |
| **Doench A375/AZD** | 5 | 2333 | Human A375/AZD cells | log2 fold-change | |
| **Doench Mouse EL4** | 4 | 951 | Mouse EL4 cells | log2 fold-change | On-target data sets with various gRNAs for mouse. Consistent quantification method. |
| **Koike-Yusa/Xu Mouse ESC** | 3 | 907 | Mouse ESC cells | log2 fold-change | |
| **Doench-CD33** | 5 | 3822 | Human MOLM13 cells | log2 fold-change | Off-target screening for human. Training sets for CFD |
| **Doench-H2-D** | 5 | 89 | Human MOLM13 cells | log2 fold-change | |
| **Hsu-EMX1 target 1,2,3,6** | 8 | 464 | Human HEK293T cells | % indels | Training sets for MIT-Zhang |
| **Tsai-EMX1/ FANCF/ VEGFA site 1-3** | 9 | 258 | Human U2OS cells | Sequencing read counts | Genome-wide off-target screening for human. |
| **Tsai-HEK293 site 1,3,4** | 9 | 150 | Human HEK293 cells | Sequencing read counts | |
| **Kuscu-sgRNA 1-12** | 10 | 2607 | Human HEK293T cells | % indels | Genome-wide off-target screening for human. |
| **Total** | | 14538 | | | |

Table S2. Summary of on-target test datasets used in this work.

| Dataset | Ref. | Number of gRNAs | Species & Biology | Efficiency Quantification | Notes |
|---|---|---|---|---|---|
| **Chari 293T** | 11 | 1234 | Human/HEK293T cells | Relative mutation rate | On-target data sets with various gRNAs for human. Frequently-used for model developing. |
| **Chari K562** | | 1239 | Human/K562 cells | Relative mutation rate | |
| **Hart Rpe** | 12 | 4214 | Human/Rpe cells | Bayes Factors | Exhausted On-target data sets with various gRNAs for human. Cell type environment considered. Good test sets for human based model developing. |
| **Hart Hct116-1 Lib 1** | | 4293 | Human/ Hct116 cells | Bayes Factors | |
| **Hart Hct116-2 Lib 1** | | 4239 | Human/ Hct116 cells | Bayes Factors | |
| **Hart HeLa Lib1** | | 4256 | Human/HeLa cells | Bayes Factors | |
| **Hart HeLa Lib2** | | 3845 | Human/HeLa cells | Bayes Factors | |
| **Varshney Zebrafish** | 13 | 102 | Zebrafish/Embryos | Relative mutation rate | On-target data sets with various gRNAs for Zebrafish. Frequently-used test sets for model developing. |
| **Gagnon Ciona** | 14 | 111 | Zebrafish/Embryos | % indels | |
| **Moreno−Mateos Zebrafish** | 15 | 1020 | Zebrafish/Embryos | % indels | |
| **Shkumatava Zebrafish** | 16 | 163 | Zebrafish/Embryos | % indels | |
| **Farboud C. elegans** | 17 | 50 | Caenorhabditis elegans | Mutagenesis rate | Alternative On-target data sets with limited gRNAs for various species. Often-used test sets for model developing. |
| **Ren Drosophila** | 18 | 39 | Drosophila | Heritable mutation rate | |
| **Liu Neuro2A Surveyor 1/0** | 19 | 205 | Mouse/Neuro2A cells | Mutagenesis rate | |
| **Eschstruth Zebrafish** | 13 | 18 | Zebrafish/Embryos | Modification frequency | |
| **Teboul Mouse In Vivo** | | 30 | Mouse/Embryos | Modification frequency | |
| **Schoenig K562 LacZ Rank** | | 24 | Human/ K562 cells | Modification frequency | |
| **Gandhi Electrop. Ciona** | 20 | 72 | Ciona/Somatic cells | Mutagenesis rate | |

Table S3. Performance of on-target activity predictions.

| Dataset | Wang Score | Doench Score | SSC | sgRNA Score | WU-CRISPR | CRISPR scan | Azimuth | uCRISPR | Kinetic model |
|---|---|---|---|---|---|---|---|---|---|
| Wang/Xu HL60 (2076) | 0.616 | 0.343 | 0.486 | 0.321 | 0.246 | 0.201 | 0.485 | 0.577 | 0.627 |
| Doench MOLM13/NB4/TF1 (881) | 0.366 | 0.497 | 0.33 | 0.273 | 0.307 | 0.01 | 0.657 | 0.606 | 0.662 |
| Doench Mouse EL4 (951) | 0.427 | 0.577 | 0.4 | 0.403 | 0.369 | 0.156 | 0.7 | 0.622 | 0.694 |
| Doench A375/AZD (2333) | 0.265 | 0.266 | 0.287 | 0.245 | 0.164 | 0.144 | 0.54 | 0.444 | 0.533 |
| Chari 293T (1234) | 0.31 | 0.246 | 0.286 | 0.457 | 0.308 | 0.123 | 0.381 | 0.518 | 0.621 |
| Koike-Yusa/Xu Mouse ESC (907) | 0.281 | 0.221 | 0.306 | 0.12 | 0.119 | 0.094 | 0.367 | 0.352 | 0.46 |
| Hart Rpe (4214) | 0.232 | 0.178 | 0.201 | 0.152 | 0.162 | 0.077 | 0.281 | 0.303 | 0.44 |
| Hart Hct116-1 Lib 1(4293) | 0.335 | 0.285 | 0.31 | 0.21 | 0.228 | 0.176 | 0.369 | 0.39 | 0.48 |
| Hart Hct116-2 Lib 1(4239) | 0.307 | 0.288 | 0.292 | 0.208 | 0.232 | 0.159 | 0.384 | 0.4 | 0.512 |
| Hart HeLa Lib1 (4256) | 0.322 | 0.253 | -0.002 | 0.192 | 0.32 | 0.144 | 0.353 | 0.375 | 0.465 |
| Hart HeLa Lib2 (3845) | 0.358 | 0.232 | 0.033 | 0.186 | 0.361 | 0.18 | 0.359 | 0.389 | 0.457 |
| Varshney Zebrafish (102) | 0.17 | 0.139 | 0.171 | 0.28 | 0.27 | 0.262 | 0.219 | 0.225 | 0.335 |
| Gagnon Ciona (111) | 0.207 | -0.072 | 0.179 | 0.202 | 0.083 | 0.357 | 0.104 | 0.239 | 0.443 |
| Moreno−Mateos Zebrafish(1020) | 0.14 | 0.038 | 0.171 | 0.145 | 0.037 | 0.579 | 0.12 | 0.166 | 0.319 |
| Shkumatava Zebrafish(163) | 0.141 | 0.088 | 0.065 | 0.128 | -0.123 | 0.251 | 0.103 | 0.142 | 0.246 |
| Wang/Xu KBM7 (2076) | 0.644 | 0.351 | 0.498 | 0.335 | 0.258 | 0.205 | 0.512 | 0.575 | 0.611 |
| Chari K562 (1239) | -0.013 | -0.019 | -0.019 | 0.009 | 0.011 | 0.013 | -0.035 | -0.044 | 0.007 |
| Farboud C. elegans (50) | 0.476 | 0.301 | 0.545 | 0.602 | 0.4 | 0.177 | 0.541 | 0.515 | 0.596 |
| Ren Drosophila (39) | 0.313 | 0.178 | 0.225 | 0.152 | -0.158 | -0.347 | 0.131 | 0.156 | 0.139 |
| Liu Neuro2A Surveyor 1/0(205) | 0.04 | 0.091 | 0.063 | 0.007 | -0.042 | -0.054 | 0.044 | 0.071 | -0.002 |
| Eschstruth Zebrafish (18) | -0.014 | 0.647 | 0.273 | 0.33 | 0.491 | -0.043 | 0.503 | 0.302 | 0.59 |
| Teboul Mouse In Vivo(30) | 0.188 | -0.083 | 0.072 | -0.154 | -0.219 | 0.426 | -0.304 | 0.016 | 0.103 |
| Schoenig K562 LacZ Rank(24) | 0.219 | 0.273 | 0.011 | 0.42 | 0.283 | -0.178 | 0.415 | 0.381 | 0.487 |
| Gandhi Electrop. Ciona(72) | 0.298 | 0.245 | 0.15 | 0.248 | 0.112 | 0.354 | 0.419 | 0.286 | 0.481 |
| Average | 0.276 | 0.232 | 0.222 | 0.228 | 0.176 | 0.144 | 0.319 | 0.332 | 0.429 |

Table S4. Performance of off-target predictions.

| Dataset | MIT_Zhang | CCTop | CROP-IT | CFD | uCRISPR | Kinetic model |
|---|---|---|---|---|---|---|
| Doench-CD33 part (3822) | 0.277 | 0.288 | 0.316 | 0.562 | 0.561 | 0.856 |
| Doench-H2-D (89) | 0.527 | 0.261 | 0.28 | 0.575 | 0.619 | 0.821 |
| Hsu-EMX1 target 1-SM (58) | 0.74 | 0.693 | 0.713 | 0.719 | 0.822 | 0.996 |
| Hsu-EMX1 target 2-SM (58) | 0.719 | 0.592 | 0.658 | 0.486 | 0.743 | 0.893 |
| Hsu-EMX1 target 3-SM (58) | 0.727 | 0.714 | 0.72 | 0.464 | 0.735 | 0.742 |
| Hsu-EMX1 target 6-SM (58) | 0.728 | 0.58 | 0.755 | 0.678 | 0.781 | 0.911 |
| Hsu-EMX1 target 1-MM (46) | 0.738 | 0.63 | 0.624 | 0.531 | 0.872 | 0.934 |
| Hsu-EMX1 target 2-MM (46) | 0.689 | 0.483 | 0.524 | 0.624 | 0.922 | 0.951 |
| Hsu-EMX1 target 3-MM (42) | 0.592 | 0.519 | 0.561 | 0.698 | 0.871 | 0.926 |
| Hsu-EMX1 target 6-MM (46) | 0.305 | 0.354 | 0.38 | 0.668 | 0.862 | 0.947 |
| Hsu-EMX1 target 1-OT (9) | 0.974 | 0.402 | 0.799 | 0.98 | 0.996 | 0.996 |
| Hsu-EMX1 target 3-OT (33) | 0.221 | 0.18 | 0.227 | 0.724 | 0.875 | 0.918 |
| Tsai-VEGFA site 1 (22) | 0.068 | 0.557 | 0.449 | 0.82 | 0.827 | 0.894 |
| Tsai-VEGFA site 3 (60) | 0.19 | 0.197 | 0.332 | 0.543 | 0.589 | 0.677 |
| Tsai-EMX1 (16) | 0.595 | 0.29 | 0.352 | 0.723 | 0.937 | 0.941 |
| Tsai-FANCF (9) | 0.367 | 0.59 | 0.375 | 0.928 | 0.97 | 0.97 |
| Tsai-HEK293 site 1 (10) | 0.619 | 0.492 | 0.417 | 0.871 | 0.916 | 0.949 |
| Tsai-HEK293 site 3 (6) | 0.138 | 0.556 | 0.625 | 0.706 | 0.985 | 0.992 |
| Tsai-VEGFA site 2 (151) | 0.499 | 0.29 | 0.299 | 0.435 | 0.39 | 0.792 |
| Tsai-HEK293 site 4 (134) | 0.404 | 0.132 | 0.259 | 0.379 | 0.452 | 0.687 |
| Wu-Nanog-sg2 (26) | 0.428 | 0.336 | 0.294 | 0.809 | 0.826 | 0.823 |
| Wu-Nanog-sg3 (5957) | 0.067 | 0.123 | 0.065 | 0.078 | 0.103 | 0.418 |
| Wu-Phc1-sg1 (2948) | 0.163 | 0.263 | 0.179 | 0.208 | 0.216 | 0.305 |
| Wu-Phc1-sg2 (663) | 0.245 | 0.168 | 0.199 | 0.272 | 0.291 | 0.392 |
| Qi-mRFP (21) | 0.45 | 0.759 | 0.769 | 0.674 | 0.777 | 0.623 |
| Kuscu-sgRNA 1 (50) | 0.698 | 0.289 | 0.327 | 0.788 | 0.806 | 0.903 |
| Kuscu-sgRNA 2 (17) | 0.533 | 0.235 | 0.466 | 0.911 | 0.963 | 0.978 |
| Kuscu-sgRNA 3 (41) | 0.462 | 0.195 | 0.195 | 0.909 | 0.927 | 0.944 |
| Kuscu-sgRNA 4 (484) | 0.072 | 0.047 | 0.077 | 0.135 | 0.187 | 0.308 |
| Kuscu-sgRNA 5 (52) | 0.089 | 0.135 | 0.343 | 0.749 | 0.819 | 0.809 |
| Kuscu-sgRNA 6 (1282) | 0.228 | 0.026 | 0.064 | 0.251 | 0.274 | 0.71 |
| Kuscu-sgRNA 7 (285) | 0.614 | 0.04 | 0.14 | 0.673 | 0.73 | 0.876 |
| Kuscu-sgRNA 8 (43) | 0.641 | 0.171 | 0.543 | 0.812 | 0.816 | 0.852 |
| Kuscu-sgRNA 9 (121) | 0.826 | 0.062 | 0.332 | 0.844 | 0.86 | 0.86 |
| Kuscu-sgRNA 10 (202) | 0.773 | 0.17 | 0.124 | 0.761 | 0.77 | 0.772 |
| Kuscu-sgRNA 11 (16) | 0.643 | 0.148 | 0.475 | 0.649 | 0.645 | 0.857 |
| Kuscu-sgRNA 12 (14) | 0.82 | 0.01 | 0.818 | 0.832 | 0.826 | 0.762 |
| Fu-EGFP Site #1-SM (20) | 0.435 | 0.847 | 0.487 | 0.71 | 0.678 | 0.687 |
| Fu-EGFP Site #2-SM (20) | 0.608 | 0.732 | 0.816 | 0.731 | 0.694 | 0.792 |
| Fu-EGFP Site #3-SM (20) | 0.357 | 0.681 | 0.543 | 0.456 | 0.448 | 0.597 |
| Fu-EGFP Site #1-MM (11) | 0.793 | 0.912 | 0.7 | 0.642 | 0.627 | 0.884 |
| Fu-EGFP Site #2-MM (11) | 0.379 | 0.527 | 0.551 | 0.866 | 0.894 | 0.879 |
| Fu-EGFP Site #3-MM (11) | 0.715 | 0.788 | 0.818 | 0.793 | 0.725 | 0.81 |

| | | | | | | |
|---|---|---|---|---|---|---|
| Fu-VEGFA Site 1 (5) | -0.367 | 0.715 | 0.83 | 0.952 | 0.897 | 0.951 |
| Fu-VEGFA Site 3 (7) | -0.29 | 0.395 | 0.391 | 0.73 | 0.724 | 0.831 |
| Cho-C4BPB (6) | 0.345 | 0.539 | 0.626 | 0.91 | 0.941 | 0.988 |
| Cho-CCR5 (14) | 0.371 | 0.585 | 0.607 | 0.85 | 0.983 | 0.978 |
| Cho-CCR5-1 (12) | 0.329 | 0.695 | 0.748 | 0.876 | 0.978 | 0.981 |
| Cho-CCR5-2 (9) | 0.436 | 0.533 | 0.582 | 0.77 | 0.993 | 0.992 |
| Cho-CCR5-3 (8) | 0.098 | 0.628 | 0.566 | 0.776 | 0.967 | 0.973 |
| Cho-CCR5-4 (9) | -0.071 | 0.645 | 0.464 | 0.774 | 0.917 | 0.929 |
| Cho-CCR5-5 (5) | -0.223 | 0.748 | 0.698 | 0.884 | 0.926 | 0.933 |
| Cho-CCR5-6 (7) | 0.448 | 0.729 | 0.319 | 0.898 | 0.991 | 0.987 |
| Cho-CCR5-7 (5) | 0.628 | 0.966 | 0.905 | 0.917 | 0.92 | 0.946 |
| Cho-CCR5-8 (6) | 0.057 | 0.692 | 0.66 | 0.693 | 0.836 | 0.892 |
| Long-DMD-mdx (33) | 0.187 | 0.14 | 0.357 | 0.694 | 0.772 | 0.805 |
| Long-DMD-WT (33) | 0.158 | 0.183 | 0.346 | 0.773 | 0.819 | 0.806 |
| Wang-sgAAVS1 (14) | 0.427 | 0.636 | 0.607 | 0.869 | 0.965 | 0.948 |
| Frock-RAG1A (33) | 0.232 | 0.199 | 0.061 | 0.239 | 0.212 | 0.233 |
| Frock-EMX1 (14) | 0.421 | 0.393 | 0.063 | 0.397 | 0.604 | 0.791 |
| Frock-VEGFA (39) | 0.231 | 0.469 | 0.405 | 0.814 | 0.841 | 0.845 |
| Kim-HBB/HAP1 (5) | 0.048 | 0.682 | 0.629 | 0.881 | 0.983 | 0.932 |
| Kim-HBB/K562 (5) | 0.046 | 0.681 | 0.628 | 0.88 | 0.982 | 0.957 |
| Kim-VEGF-A/HAP1 (9) | -0.199 | 0.538 | 0.446 | 0.746 | 0.924 | 0.979 |
| Kim-VEGF-A/K562 (9) | -0.177 | 0.578 | 0.51 | 0.817 | 0.929 | 0.934 |
| Ran-EMX1 target1 (12) | 0.471 | 0.15 | 0.527 | 0.823 | 0.811 | 0.831 |
| Ran-EMX1 target2 (9) | -0.073 | 0.48 | 0.376 | 0.913 | 0.977 | 0.976 |
| Kleinstiver-EMX1 site 1 (9) | 0.55 | 0.413 | 0.338 | 0.709 | 0.929 | 0.942 |
| Kleinstiver-EMX1 site 2 (11) | 0.903 | 0.147 | 0.463 | 0.856 | 0.944 | 0.943 |
| Kleinstiver-FANCF site 2 (26) | 0.622 | 0.266 | 0.342 | 0.527 | 0.839 | 0.912 |
| Kleinstiver-FANCF site 3 (9) | 0.741 | 0.603 | 0.295 | 0.792 | 0.949 | 0.987 |
| Kleinstiver-ZSCAN2 (9) | 0.267 | 0.079 | 0.497 | 0.924 | 0.977 | 0.978 |
| Kim-VEGFA1 (26) | 0.039 | 0.682 | 0.586 | 0.815 | 0.798 | 0.814 |
| Kim-VEGFA2 (22) | 0.002 | 0.424 | 0.383 | 0.22 | 0.637 | 0.871 |
| Kim-VEGFA3 (25) | -0.074 | 0.412 | 0.404 | 0.559 | 0.465 | 0.764 |
| Kim-EMX1 (12) | 0.507 | 0.485 | 0.121 | 0.674 | 0.842 | 0.932 |
| Kim-FANCF (9) | 0.425 | 0.545 | 0.357 | 0.888 | 0.907 | 0.953 |
| Kim-HEK293-1 (8) | 0.777 | 0.27 | 0.563 | 0.589 | 0.687 | 0.787 |
| Kim-HEK293-3 (8) | 0.395 | 0.195 | 0.511 | 0.656 | 0.931 | 0.966 |
| Kim-HEK293-4 (20) | -0.071 | 0.421 | 0.487 | 0.637 | 0.841 | 0.894 |
| Slaymaker-VEGFA(1)-SM (58) | 0.713 | 0.63 | 0.795 | 0.746 | 0.749 | 0.883 |
| Slaymaker-VEGFA(1)-MM (19) | 0.562 | 0.7 | 0.807 | 0.864 | 0.846 | 0.942 |
| Slaymaker-VEGFA(1)-OT (11) | 0.153 | 0.267 | 0.503 | 0.827 | 0.801 | 0.901 |
| Liu-Renilla (21) | 0.603 | 0.864 | 0.781 | 0.355 | 0.659 | 0.762 |
| Average | 0.378 | 0.438 | 0.471 | 0.692 | 0.747 | 0.848 |
